# Human motor units in microfluidic devices are impaired by FUS mutations and improved by HDAC6 inhibition

**DOI:** 10.1101/2020.10.21.346874

**Authors:** Katarina Stoklund Dittlau, Emily N. Krasnow, Laura Fumagalli, Tijs Vandoorne, Pieter Baatsen, Axelle Kerstens, Giorgia Giacomazzi, Benjamin Pavie, Maurilio Sampaolesi, Philip Van Damme, Ludo Van Den Bosch

## Abstract

Neuromuscular junctions (NMJs) ensure proper communication between motor neurons and muscle through the release of neurotransmitters. In motor neuron disorders, such as amyotrophic lateral sclerosis (ALS), NMJs degenerate resulting in muscle atrophy, paralysis and respiratory failure. The aim of this study was to establish a versatile and reproducible *in vitro* model of a human motor unit to study the effect of ALS-causing mutations. Therefore, we generated a co-culture of human induced pluripotent stem cell-derived motor neurons and human primary mesoangioblast-derived myotubes in microfluidic devices. A chemotactic and volumetric gradient facilitated the growth of motor neuron neurites through microgrooves resulting in the interaction with myotubes and the formation of NMJs. We observed that ALS-causing *FUS* mutations resulted in a reduced neurite outgrowth and in a decreased NMJ number. Interestingly, the selective HDAC6 inhibitor, Tubastatin A, improved the neurite outgrowth and the NMJ morphology of *FUS*-ALS co-cultures, further prompting HDAC6 inhibition as a potential therapeutic strategy for ALS.

## Introduction

Neuromuscular junctions (NMJs) are crucial for the communication between spinal lower motor neurons (MNs) and skeletal muscle fibers (1). These NMJs are vulnerable in neurodegenerative diseases, such as amyotrophic lateral sclerosis (ALS) and spinal muscular atrophy (SMA), where the degeneration of NMJs results in muscle weakness and atrophy, ultimately causing respiratory insufficiency and patient death (2–5). There is ample evidence that the disconnection of the motor axon from the muscle is the first step in the disease process of these motor neuron disorders (6–9) making it an important mechanism to study. Initially, the motor neuron will compensate for this retraction by axonal sprouting and collateral re-innervation. However, when the disease progresses, these compensatory mechanisms fail and the motor neuron eventually dies, a phenomenon defined as the ‘dying-back’ mechanism (7,10). Symptoms in patients start to occur when large populations of motor neurons are affected, resulting in atrophy and weakness of muscle groups. Motor neurons have very long axons, which makes them more susceptible to this ‘dying-back’ mechanism compared to other neurons, partially explaining the selective vulnerability in ALS and SMA pathogenesis (6–13). Familial ALS constitutes approximately 10% of patients, and is mainly caused by mutations in the *chromosome 9 open reading frame 72* (*C9ORF72*), *superoxide dismutase 1* (*SOD1*), *TAR DNA-binding protein 43* (*TARDBP*), or *fused in sarcoma* (*FUS*) genes, while the remaining 90% of cases are sporadic with unknown aetiology (14–18). Evidence of ‘dying back’ and loss of NMJs has been reported in several ALS-mouse models and in patients (8,9,12,19,20). However, there are currently limited versatile *in vitro* models available to investigate this phenomenon in a human context.

In order to further study the NMJs and their relation to neurodegenerative diseases, it is important to have a standardized, human cell-derived model of these NMJs *in vitro*, for which co-culturing of MNs and skeletal muscle is required. Co-cultures of these highly specialized cell types in single compartments have been developed using both human and animal cells, and in some studies, functional NMJs were established (21–26). However, such models are unable to account for the unique culture microenvironments occupied by cell-specific domains of MNs and muscle fibers (27). The use of “Campenot” chambers (28) and microfluidic devices offers the opportunity to overcome this problem (29–36). The fluidically isolated compartments, between which only neurites can grow, not only allows for maintenance of cell-type specific microenvironments, but also allows for isolation of subcellular compartments, such as the distal and the proximal part of the axon for region-specific analyses (29,31).

While the development of customized microfluidic platforms (36,37) has significantly improved the potential for disease modelling, a standardized accessible method for the formation of human NMJs *in vitro* is lacking. Therefore, we aimed to develop a human-derived system, which combines commercially available microfluidic devices, iPSC-derived MNs, and mesoangioblast-derived myotubes to generate and study functional human NMJs in a compartmentalized system. With the supplementation of agrin and laminin, we significantly increased the clustering of nicotinic acetylcholine receptors (AChRs), the outgrowth of neurites and the formation of NMJs. Using this method, we discovered a compromised outgrowth of MN neurites and an impairment in NMJ numbers in *FUS*-mutant co-cultures in comparison to their CRISPR/Cas9 gene-edited isogenic controls. Interestingly, we could partially restore this phenotype by treating both compartments with the selective histone deacetylase 6 (HDAC6) inhibitor, Tubastatin A. Our findings show that this *in vitro* model is a valuable tool to assess human NMJ physiology in health and disease.

## Results

### Generation of human iPSC-derived motor neurons and mesoangioblast-derived myotubes

To generate human NMJs *in vitro*, we made use of human iPSCs (hiPSCs) and human primary mesoangioblasts (MABs). MABs are vessel-associated mesenchymal stem cells, which are isolated from adult skeletal muscle tissue (38–40). MABs were expanded and subsequently differentiated into myotubes over a 10-day period, while hiPSCs were differentiated into MNs via a 28-day protocol (41). MN differentiation efficiency and MAB fusion index were evaluated using immunocytochemistry (ICC). At day 28 of MN differentiation, the hiPSC-derived MNs stained positive for MN-specific markers neurofilament heavy chain (NEFH) (84.7 ± 1.9%), choline acetyltransferase (ChAT) (75.0 ± 2.9%) and Islet-1 (93.8 ± 1.7%) in addition to the pan-neuronal marker βIII-tubulin (90.8 ± 1.5%) (fig. 1a, b). These values are similar to previous studies utilizing the same MN differentiation protocol (41,42). After 10 days of differentiation, the human MAB-derived multinucleated myotubes stained positive for muscle markers desmin (8.7 ± 1.3%), myosin heavy chain (MyHC) (8.4 ± 1,2%), myogenin (MyoG) (5.8 ± 0,8%) and titin (7.4 ± 0,8%) (fig. 1c,d; supplementary fig. 1).

**Fig. 1:**
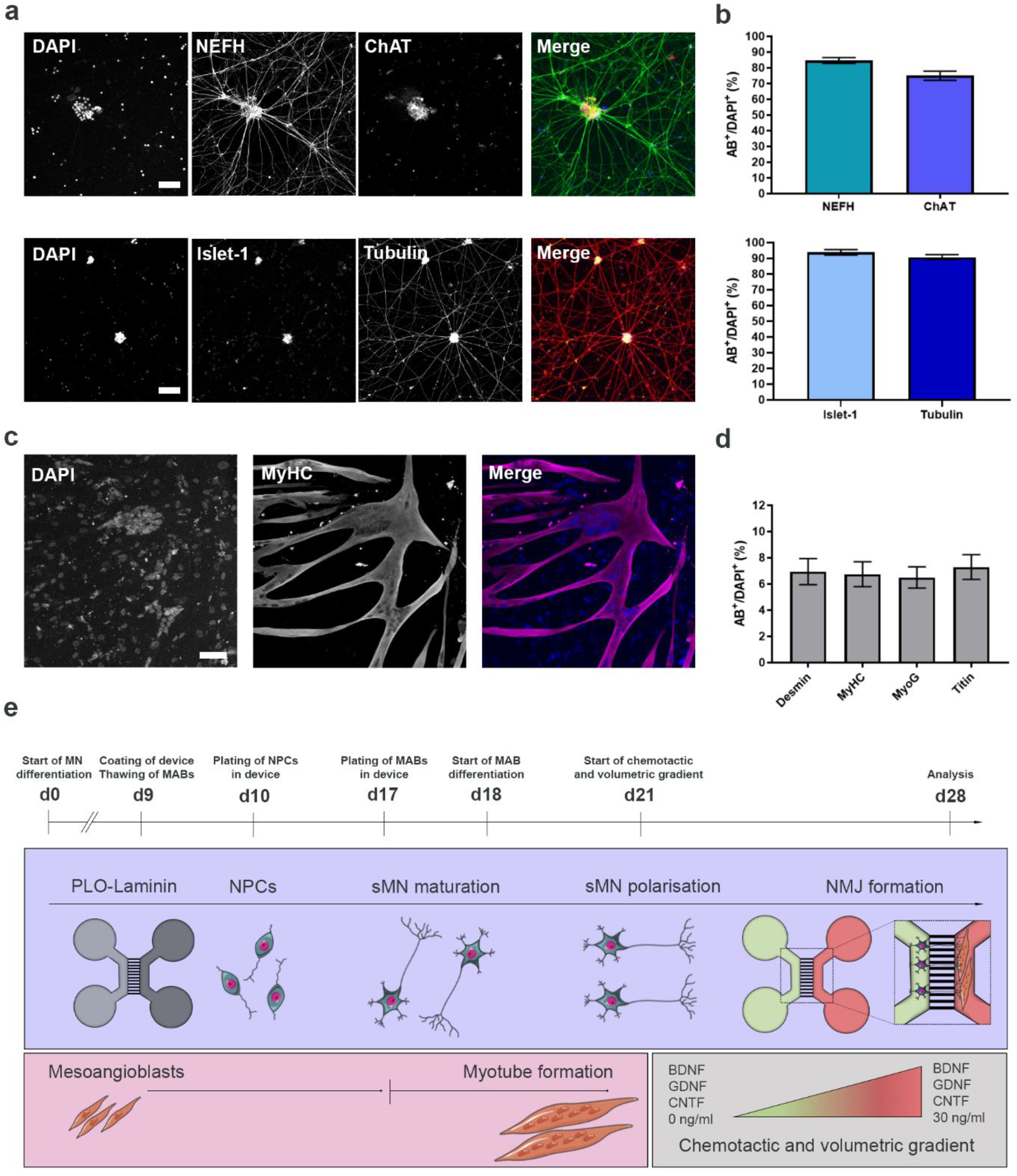
Characterisation of monocultures and overview of NMJ protocol. **a.** Confocal images of cells stained with motor neuron (MN) markers neurofilament heavy chain (NEFH), choline acetyltransferase (ChAT) and Islet-1 as well as pan-neuronal marker βIII-tubulin at day 28 of MN differentiation. Scale bar: 75 μm. **b.** Number of cells positive for MN and pan-neuronal markers. **c.** Confocal images of myotube heavy chain (MyHC)-positive myotubes 10 days after initiation of differentiation. Scale bar: 75 μm. **d.** Quantification of mesoangioblast (MAB) fusion index with myotube markers Desmin, MyHC, Myogenin (MyoG) and titin. **e.** Schematic overview of co-culture protocol and differentiation timeline. At day 0 (d0), differentiation of iPSCs into MN is started. On day 9 (d9), microfluidic devices are coated with poly-L-ornithine (PLO) and laminin, and MABs are thawed for expansion in T75/T175 flasks. On day 10 (d10), motor neural progenitor cells (NPCs) are plated on one side (light grey compartment) of the pre-coated device. One week later (d17), MABs are seeded in the opposite side of the device (dark grey compartment). Myotube differentiation is initiated (d18). On day 21 (d21), a volumetric and chemotactic gradient of neurotrophic factors (BDNF, GDNF and CNTF) is implemented to facilitate polarized growth of spinal motor neurons (sMN) through the microgrooves towards the myotube compartment to initiate the formation of neuromuscular junctions (NMJ). Cell illustrations are modified from Smart Servier Medical Art licensed under a Creative Commons Attribution 3.0 Unported License (https://creativecommons.org/licenses/by/3.0/). The data in **b** (n=15) and **d** (n=15) represent mean ± s.e.m from three independent experiments.

### NMJs spontaneously form in a motor neuron-myotube microfluidic co-culture system

In order to generate NMJs through co-culturing of MNs and MABs, we utilized two types of Xona™ microfluidic devices (SND75 and XC150). To ensure compartmentalization, motor neuron neural progenitor cells (MN-NPCs) were plated at day 10 of MN differentiation on one side of the device (fig. 1e: light grey compartment) and allowed to differentiate for one week, before MABs were plated at day 17 of MN differentiation on the other side of the fluidically isolated device (fig. 1e: dark grey compartment, supplementary fig. 2). The difference in plating time was conducted to ensure an efficient co-culture between maturing MNs and fusing myotubes, in order to make full use of the culture window in which MAB-derived myotubes are viable.

On day 21 of MN differentiation, a chemotactic and volumetric gradient was implemented to induce migration of MN axons (fig. 1e, green compartment) through the microgrooves to the compartment of the device containing myotubes (fig. 1e, red compartment). The hydrostatic pressure created by the different volumes led to fluidic isolation of the myotube compartment from the MN compartment, while the 300% growth factor gradient of glial cell line-derived neurotrophic factor (GDNF), brain-derived neurotrophic factor (BDNF) and ciliary neurotrophic factor (CNTF) promoted the polarized growth of the MNs. Due to these gradients, a larger proportion of axons migrated through the microgrooves from the MN compartment towards the myotube compartment (fig. 2a), where they were able to connect with clusters of α-bungarotoxin (Btx)-positive AChRs expressed on myotubes. AChRs represent an important guidance cue for MN axons and hence NMJ formation *in vivo* (43) and we could indeed observe interactions of MN neurites and AChR clusters on myotubes indicative of the formation of NMJ-like structures (referred to as NMJs from here on) (fig. 2.b, c). However, only a limited number of AChR clusters and hence NMJs were identified in each culture.

**Fig. 2:**
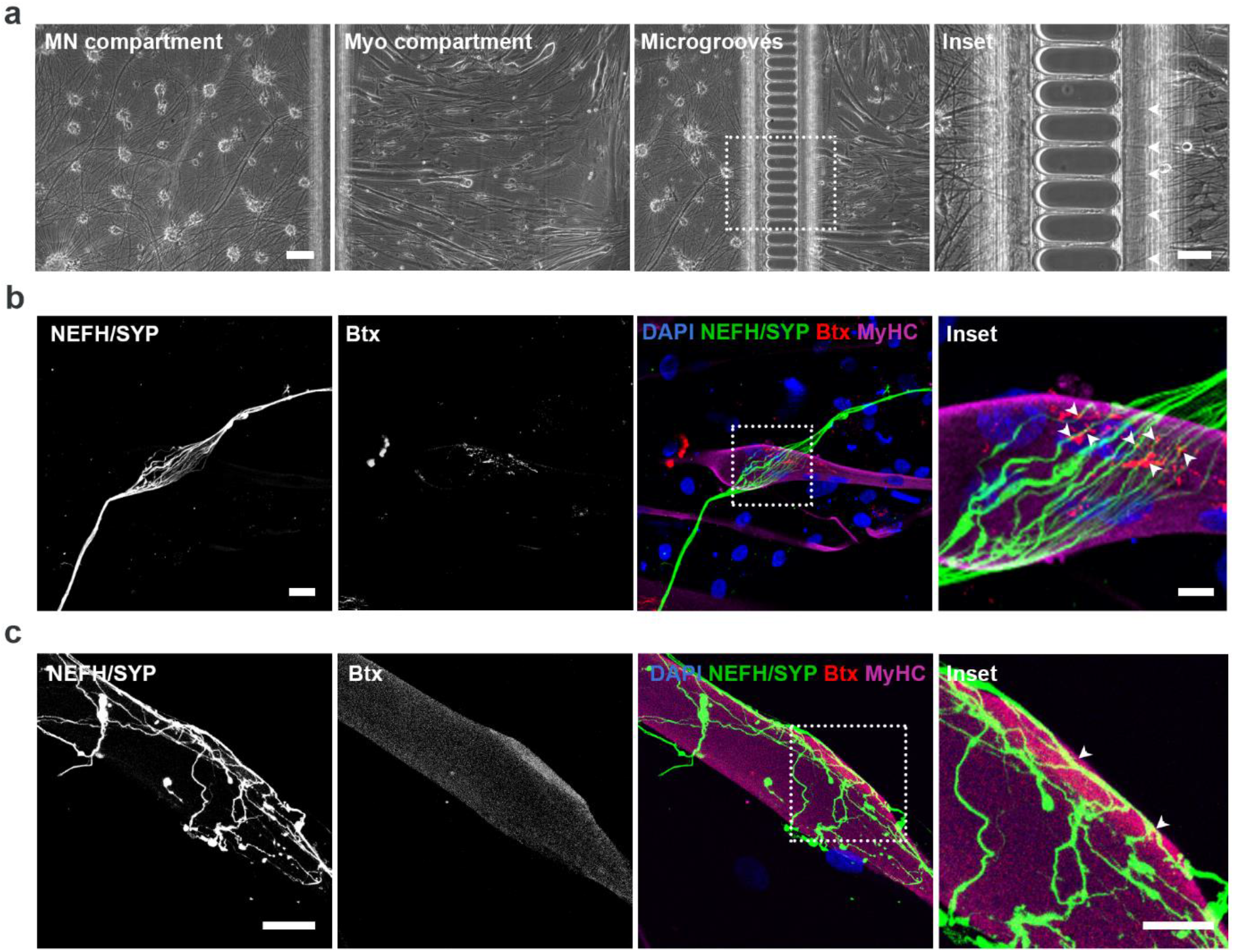
Co-culturing of MNs and myotubes in microfluidic devices leads to NMJ formation. **a.** Bright field micrographs of MN and myotube (Myo) compartments in XC150 microfluidic device at day 28 of MN differentiation and day 10 of myotube differentiation. Scale bar: 100 μm. Inset represents a magnification of axonal migration through microgrooves (arrowheads). Inset scale bar: 50 μm. **b.** Confocal micrograph of neuromuscular junction formation are shown as co-localization of presynaptic marker synaptophysin (SYP) and acetylcholine receptor marker α-bungarotoxin (Btx) on MyHC-labelled myotubes. Scale bar: 25 μm. Inset displays a magnification of co-localizations (arrowheads). Inset scale bar: 10 μm. **c.** Confocal micrograph magnification of neuromuscular junction. Scale bar: 25 μm. Inset shows a magnification of co-localizations of SYP and Btx (arrowheads). Inset scale bar: 10 μm. The data represent four independent experiments.

### Agrin and laminin stimulate neurite outgrowth and NMJ formation

In order to increase the NMJ numbers in our model, we tested whether supplementing the myotube culture medium with agrin and laminin (A/L) had an effect, as it has been demonstrated that these compounds can increase the AChR clustering at the sarcolemma (43–45). Previous reports demonstrated how AChR clustering on cultured myotubes is independent of neuronal presence (46,47), so to investigate the effect on human primary MABs-derived myotubes, we quantified AChR clustering with and without supplementation of these compounds (supplementary fig. 3). This revealed an increase in AChR cluster numbers per myotube with agrin and laminin supplements (2.8 ± 0.3) in comparison to untreated controls (1.9 ± 0.2, supplementary fig. 3c). Agrin and laminin supplements did not affect the MABs ability to fuse into myotubes (supplementary fig. 1; A/L: Desmin: 6.9 ± 1.0%; MyHC: 6.7 ± 0.9; MyoG: 6.5 ± 0.8%, titin: 7.3 ± 0.9%).

To quantify the number of human NMJs generated in our co-culture system and to characterize the NMJ morphologies, we utilized ICC. NMJs were identified through co-localisation of distal neurites (NEFH and synaptophysin (SYP)) with Btx-positive AChRs on MyHC-labelled myotubes and the number of interactions was normalized to the number of myotubes. Two types of NMJ morphologies were observed. They either appeared rudimentary and circular as single contact point NMJs (fig. 3a, supplementary fig. 4a), or elongated with broad, flat, irregular multiple contact point NMJs (fig. 3b, supplementary fig. 4b). In most cases, contacts between axons and myotubes did not produce distinct axon termination in endplate formation as seen *in vivo* but revealed a continuation of axonal growth post-embedment into myotubes (fig. 3b). No obvious morphological differences in size, shape or embedment-form were observed immunocytochemically between controls and conditions with agrin and laminin (fig. 3, supplementary fig. 4). However, the addition of agrin and laminin doubled the total number of NMJs per myotube, assessed by co-localisation between SYP-Btx (4.9 ± 0.6) in comparison to control devices without agrin and laminin supplementation (2.5 ± 0.4, fig. 3c). The addition of agrin and laminin likewise increased the number of multiple contact point NMJs (3.0 ± 0.4) in comparison to single contact point NMJs (1.9 ± 0.3, fig. 3d). A difference, which was not seen in the control devices (multiple: 1.4 ± 0.2; single: 1.0 ± 0.2). To further investigate the morphology of MN-myotube co-cultures and human NMJs *in vitro*, we performed scanning electron microscopy (SEM). SEM images revealed neurite embedment into the surface of myotubes (fig. 3e, f) in line with our ICC observations. Here, we also identified two types of MN-myotube interactions: simple, single contact point NMJs (fig. 3e) and larger, complex, multiple contact point NMJs (fig. 3f). In addition, we observed larger, elongated NMJs in agrin and laminin supplemented conditions compared to untreated controls further confirming the beneficial effects of these compounds.

**Fig. 3:**
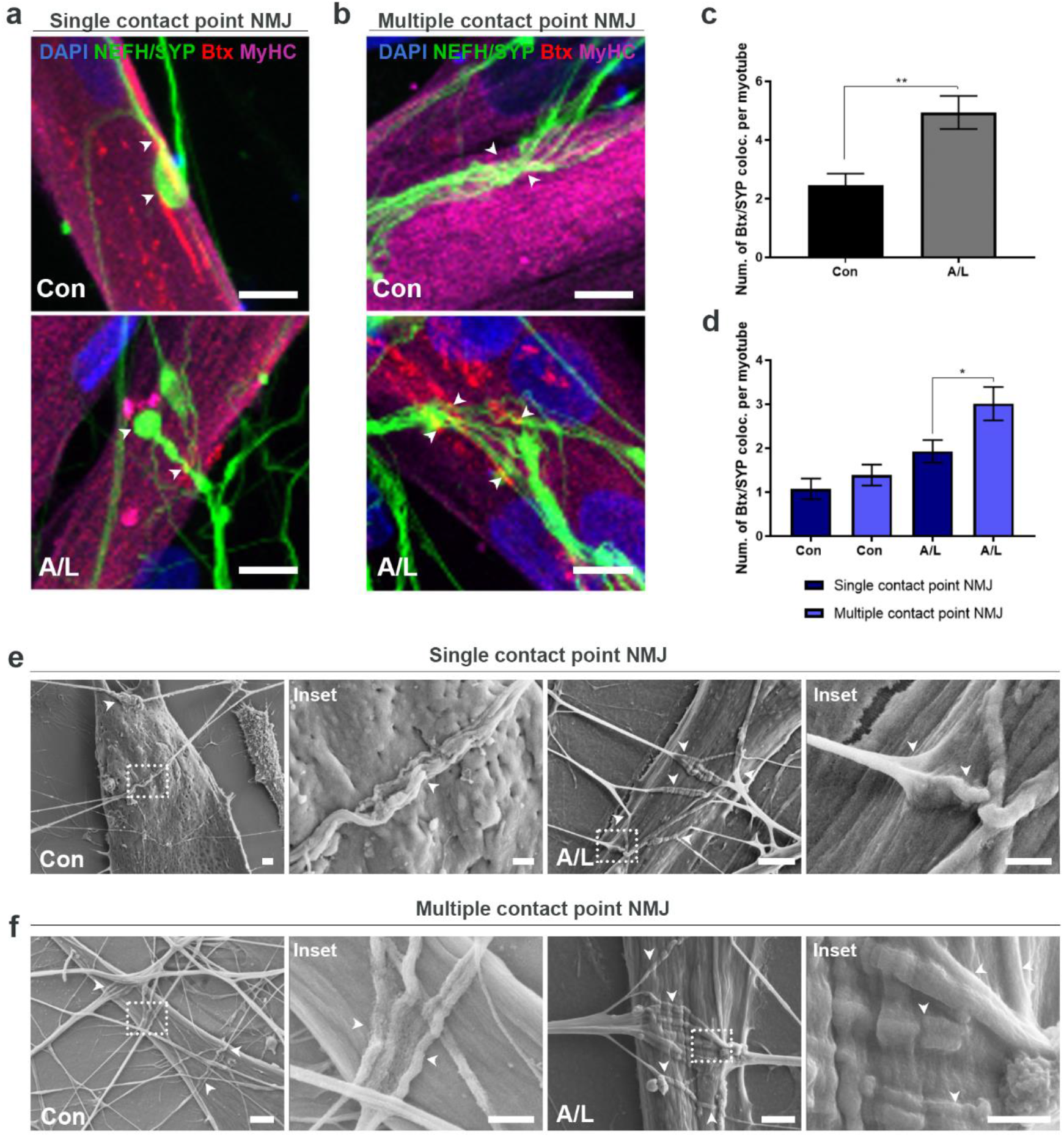
Agrin and laminin improve NMJ formation in microfluidic devices. **a-b.** Confocal micrographs of NMJs in agrin (0.01 μg/ml) and laminin (20 μg/ml) supplemented conditions (A/L) and untreated controls (Con) at 28 days of motor neuron differentiation in XC150 microfluidic devices. NEFH- and SYP-positive motor neuronal axons either form a single contact point connection with myotubes forming rudimental NMJs (**a**) or they fan out upon interaction with myotubes and create multiple contact point NMJs (**b**). Scale bar: 10 μm. Arrowheads mark colocalizations between SYP and Btx. **c.** Number of Btx and SYP co-localisations per myotube. **d.** Quantifications of NMJ single and multiple contact point morphology. **e-f.** Representative scanning electron microscopy images of NMJ formation in SND75 microfluidic devices at day 28 of motor neuron differentiation in A/L supplemented and control conditions. NMJs appear either as rudimentary single contact point (**e**) or as larger connections through multiple contact points (**f**) with myotubes. Scale bar: 2 μm. Insets represent magnifications of NMJ formations (arrowheads). Inset scale bar: 1 μm. The data (n=34-50) in **c** and **d** represent mean ± s.e.m from four independent experiments. Statistical analysis in panel **c** was performed using unpaired t-test with significant difference indicated by **p<0.01. One-way Anova with Tukey’s multiple comparisons test was performed in panel **d** with significant difference indicated by *p<0.05.

As agrin and laminin are signalling molecules important for neurite guidance, we investigated whether these supplements have an effect on neurite outgrowth in addition to NMJ formation. In order to quantify the difference, we acquired tile scan images of the entire myotube compartment and isolated the neurites (supplementary fig. 5). The images were converted to a mask and analysed with a linear Scholl analysis script similar to a previously published method (48) (supplementary fig. 5a, b). The Scholl analysis creates an intersection line every 50 μm, and the number of pixels per intersection was quantified (supplementary fig. 5c). Due to the high density of neurites, which grow through the microgrooves in thick bundles, we omitted the first intersection at 50 μm distance from the microgroove exit. The initial exponential increase in pixel intersections correlated with the ‘debundling’ and spreading of neurites post microgroove crossing (supplementary fig. 5c). Interestingly, the linear Scholl quantifications revealed an increase in neurite outgrowth due to agrin and laminin treatment (supplementary fig. 5c).

Taken together, these results demonstrate the formation of human NMJs in our *in vitro* system. Supplementation of agrin and laminin promoted the outgrowth of neurites and increased the total number of NMJs per myotube as well as the contact areas between branching motor neuron axons and myotubes further enhancing the embedment.

### Human *in vitro* NMJs are functional through MN stimulation

To evaluate whether the human NMJs were functional, we performed live-cell Ca^2+^ imaging (fig. 4). MN somas and proximal axons were depolarized using KCl and a subsequent increase in Ca^2+^ influx in the myotubes preloaded with the Ca^2+^-sensitive Fluo-4 dye was measured (fig. 4a). The fluidic isolation of the compartments in the microfluidic device ensures no direct contact between the high KCl solution and the myotubes as previously confirmed (supplementary fig. S2a). Stimulation of MNs resulted in a single Ca^2+^ transient wave in the MN-innervated myotubes indicating a functional connection through NMJ formation (fig. 4b, c).

**Fig. 4:**
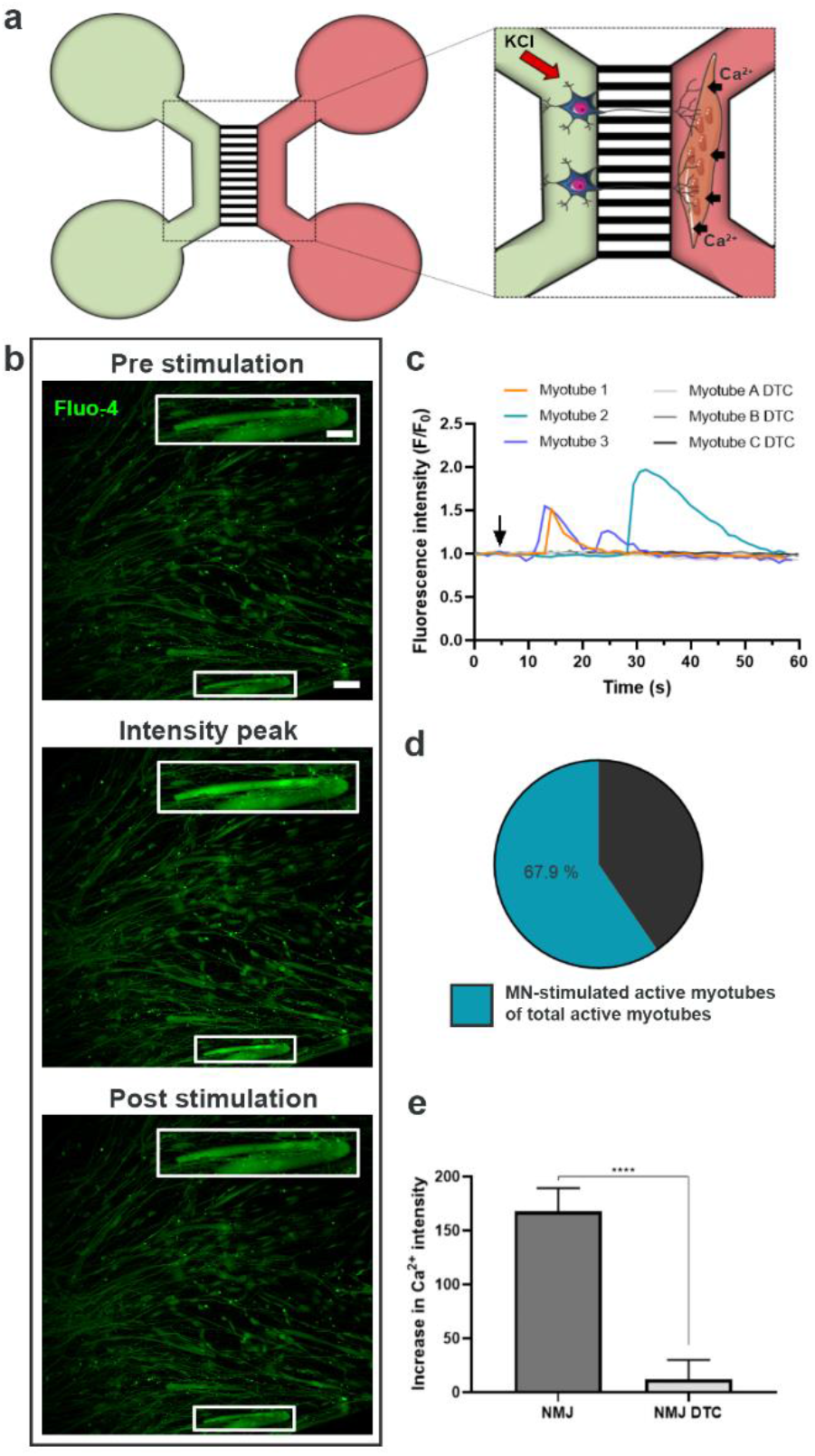
*In vitro* NMJs are functionally active. **a.** Schematic overview of transient fluorescent calcium (Ca^2+^) imaging in XC150 microfluidic devices at day 28 of motor neuron (MN) differentiation. MN somas and proximal axons are stimulated with 50 mM KCl on one side of the fluidically isolated microfluidic device (green compartment). The stimulation initiates an intracellular response through MN axons, towards the myotube compartment (red compartment), which evokes an increase in Ca^2+^ influx in the Fluo-4-calcium-loaded myotubes. Cell illustrations are modified from Smart Servier Medical Art licensed under a Creative Commons Attribution 3.0 Unported License (https://creativecommons.org/licenses/by/3.0/). **b.** Fluorescent micrograph examples of prior, during and post-stimulation depicting a wave of increase in intracellular Ca^2+^ in myotubes upon MN stimulation. Scalebar: 100 μm. Insets magnifies an innervated active myotube. Inset scale bar: 200 μm. **c.** Representative Ca^2+^ influx curves in myotubes after KCl activation (arrow) of MNs before (Myotube 1-3) and after 10 min treatment with 19 μM nicotinic AChR competitive antagonist tubocurarine (Myotube A-C DTC). **d.** Ratio of MN-stimulated active myotube to directly KCl-activated myotubes (n=38). **e.** Intensity fluorescent increase due to Ca^2+^ influx (n=31). Statistical analysis in panel **e** was performed using Mann-Whitney test with ****p<0.0001. The data represent mean ± s.e.m from three-four independent experiments.

In order to differentiate between excitable and non-excitable myotubes, we also stimulated the myotubes directly with KCl in the myotube compartment post MN-stimulation (supplementary fig. 6a). Excitable myotubes were considered responsive to KCl through characteristic single Ca^2+^ wave formation. Approximately 68% (67.9 ± 11.8%) of myotubes of total directly KCl-stimulated active myotubes were responsive through MN stimulation (fig. 4d). Quantifications of peak value of the Ca^2+^ influx were performed on myotubes demonstrating MN-dependent activity (fig. 4e). To confirm that the Ca^2+^ transient was due to NMJ transmission, NMJs were blocked using the nicotinic AChR competitive antagonist, tubocurarine (DTC) (49). This AChR-specific blocker successfully inhibited the Ca^2+^ transients in our system (fig. 4c, e). To assess whether the myotube excitability was dependent on MN presence, we cultured myotubes separately and stimulated them with KCl (supplementary fig. 6b, c). Myotubes were active regardless of MN presence (supplementary fig. 6b), and they likewise emitted a Ca^2+^-response independent of DTC treatment, excluding a direct inhibitory effect of DTC on the muscle membrane potential (supplementary fig. 6c). In conclusion, these results confirm that the observed Ca^2+^-peaks in the myotubes after depolarisation of the MNs resulted from the NMJ-mediated myotube activation.

### Mutant *FUS* causes impaired neurite outgrowth and a reduction in NMJ numbers

To assess the effect of disease-causing ALS mutations in this human *in vitro* NMJ model, we used two mutant *FUS* iPSC lines, one from a 17-year old ALS patient with a *de novo* point mutation (P525L) and another one from a 71-year old ALS patient with a R521H mutation. These patient lines were systematically compared to their corresponding CRISPR/Cas9 gene-edited isogenic P525P and R521R controls, respectively (41). MN differentiation verification with ICC showed no difference in the differentiation potential between the different lines and revealed an approximate expression of 85-95% of MN markers (supplementary fig. 7).

ICC analysis of the NMJs discovered a significant difference in the total number of NMJs per myotube between the P525P isogenic control (4.8 ± 0.4) and the MNs containing the aggressive P525L mutation in *FUS* (2.7 ± 0.3, fig. 5a-b, supplementary fig. 8) indicating a mutant *FUS*-dependent impairment of NMJs. A small but significant difference was also observed between the R521R (4.5 ± 0.3) and R521H (3.7 ± 0.5) MN/myotube co-culture systems. In addition, we observed a significantly higher number of multiple contact point NMJs (P525P: 3.5 ± 0.4, R521R: 3.2 ± 0.3) than single contact point NMJs (P525P: 1.3 ± 0.1, R521R: 1.2 ± 0.2, fig. 5c) in the isogenic control systems, while this difference was not observed in the mutated systems (P525L: multiple: 1.0 ± 0.2; single: 1.6 ± 0.2, R521H: multiple: 2.0 ± 0.3; single: 1.8 ± 0.3) suggesting that there is a difference in the maturation state between the ALS and control co-cultures.

**Fig. 5:**
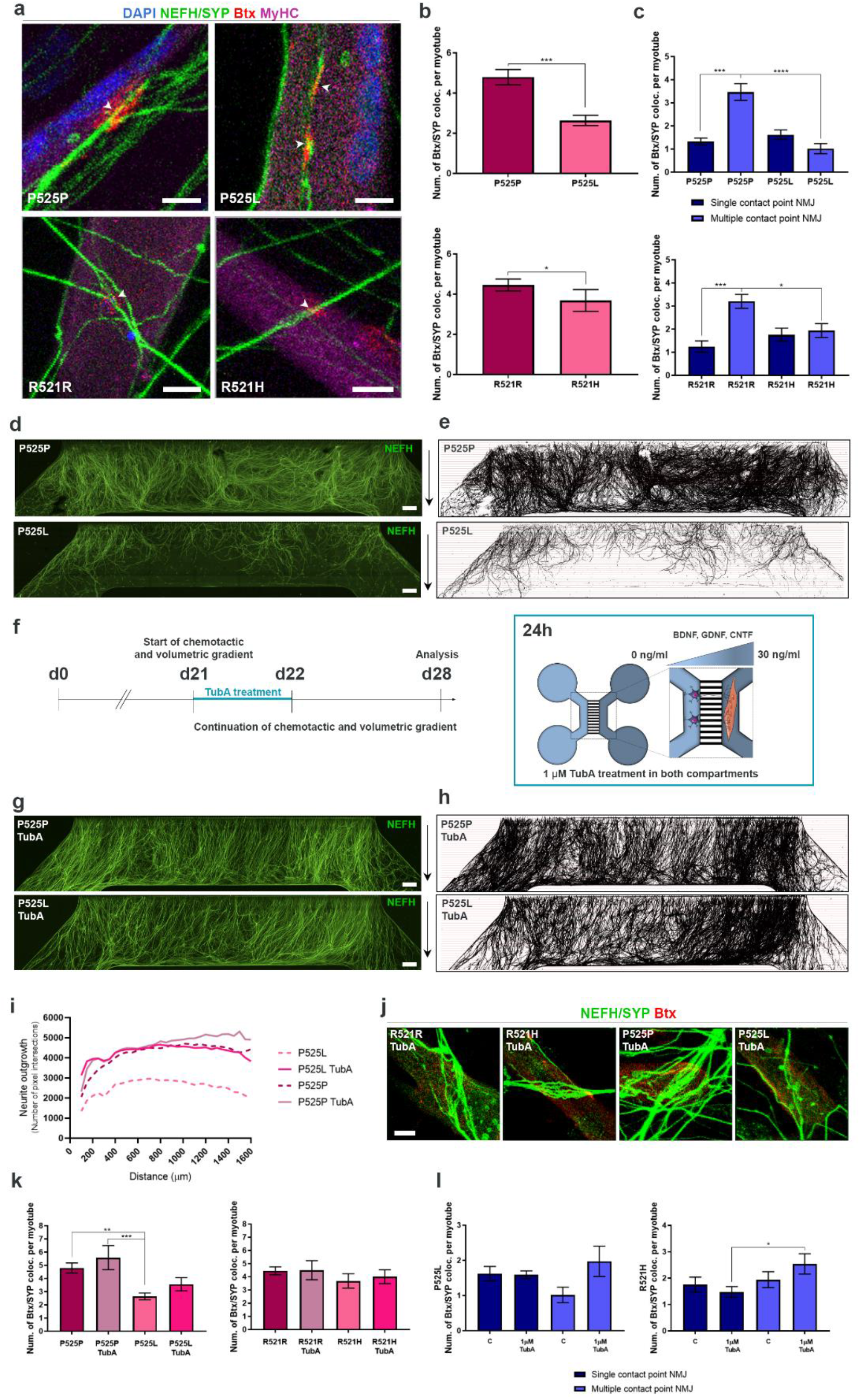
ALS-dependent impairment of NMJ numbers and neurite outgrowth is improved by HDAC6 inhibition. **a.** Confocal micrographs of NMJs in agrin and laminin supplemented conditions from mutant *FUS* MN/myotube co-cultures (P525L, R521H) and isogenic control MN/myotube co-cultures (P525P, R521R) at 28 days of motor neuron differentiation in XC150 microfluidic devices. Scale bar: 10 μm. Arrowheads mark the co-localizations between SYP and Btx. **b.** Quantification of NMJs as Btx and SYP co-localisations and expressed per myotube. **c.** Quantification of NMJ single and multiple contact point morphology. **d.** Tile scan confocal overviews of neurite outgrowth (NEFH) in myotube compartment from P525L and P525P cultures. Arrows (right) depict growth direction from exit of microgrooves. Scale bar: 300 μm. **e.** Masks of tile scans with intersection lines at every 50 μm starting from microgroove exit. **f.** Schematic representation of the experimental time course implementing treatment with Tubastatin A (TubA). At day 21 (d21), both MN and myotube compartments were treated with 1μM TubA for 24 h in addition to the start of the chemotactic and volumetric gradient. Cell illustrations are modified from Smart Servier Medical Art licensed under a Creative Commons Attribution 3.0 Unported License (https://creativecommons.org/licenses/by/3.0/). **g.** Tile scan confocal overviews of neurite outgrowth in myotube compartment from P525L and P525P MN/myotube co-cultures after 24 h of TubA treatment. Scale bar: 300 μm. **h.** Masks of tile scans after 24 h of TubA treatment. **i.** Neurite outgrowth quantifications of the number of pixel intersections in P525L and P525P MN/myotube co-cultures before and after a 24 h treatment with TubA. **j.** Confocal micrographs of NMJs after 24 h of TubA treatment. Scale bar: 10 μm. **k.** Quantification of NMJs as Btx and SYP co-localisation per myotube before and after TubA treatment. Control data before TubA treatment is identical to panel b. **l.** Morphological analysis of NMJs before (c) and after TubA treatment. Control data (c) are identical to panel c. All data are from three to eight independent experiments. The data in **b, c, k** and **l** represent mean ± s.e.m. Statistical analysis in panel **b** (n=20-34) was performed using unpaired t-test and Mann-Whitney test. One-way Anova with Tukey’s multiple comparisons test and Kruskal-Wallis test with Dunn’s multiple comparisons test were used in panel **c** (n=20-34), **k** (n=16-37) and **l** (n=16-37). Significant differences are indicated by *p<0.05, **p<0.01, ***p<0.001 and ****p<0.0001.

Interestingly, we also observed a striking difference in neurite outgrowth from the MN compartment towards the myotube compartment between the P525L mutant in *FUS* and its isogenic control (P525P) (fig. 5d, e). The linear Scholl analysis confirmed this observation and revealed a decrease in neurite outgrowth in the mutant (P525L) system compared to the isogenic control (P525P) MNs (fig. 5i). No obvious difference in outgrowth was observed in the *FUS*-mutant (R521H) and isogenic control (R521R) MN/myotube co-cultures (supplementary fig. 9a, b, e). Next, we investigated whether this difference in neurite outgrowth was dependent on myotube presence (supplementary fig. 9f-k). Interestingly, we observed fewer neurites crossing in smaller bundles in the devices without the myotubes, suggesting that myotubes play a role in the polarization of neurites in addition to the volumetric and chemotactic gradient. A similar decrease in neurite outgrowth in *FUS*-mutant P525L in comparison to isogenic control P525P systems was observed without myotube presence although the difference was less pronounced. To test whether mutant and isogenic control MNs showed any variation in their initial neurite outgrowth, we cultured MN-NPCs from mutant *FUS* (P525L) and its isogenic control (P525P) in 24-well plates and analysed the neurite outgrowth using IncuCyte NeuroTrack software (supplementary fig. 10). Images were taken every 2 h over the first 24 h after plating of NPCs. No difference in neurite outgrowth was observed between the two lines within the first 24 h, suggesting that this phenotype only appears later upon MN maturation.

Taken together, these results showed that the number of NMJs and the neurite outgrowth of motor neurons can be affected by mutations in *FUS* in our human co-culture system. These impairments can be successfully rescued by correction of the mutation, emphasising that the effects are a direct cause of the point mutation in *FUS*.

### HDAC6 inhibition improves mutant *FUS*-mediated impairments of neurite outgrowth and NMJ formation

Many studies demonstrated beneficial effects of HDAC6 inhibition in various models of neurodegenerative diseases (41,50–55). HDAC6 is primarily found in the cytosol where it can deacetylate α-tubulin, which affects axonal transport (reviewed in (56,57)). We previously showed that HDAC6 inhibition can rescue axonal transport defects in mutant *FUS* motor neurons (41). As the impairment in neurite outgrowth could correlate with axonal transport deficits, we investigated whether HDAC6 inhibition could also improve this phenotype.

Therefore, we treated our co-cultures with the specific HDAC6 inhibitor, Tubastatin A (TubA; 1μM), in both the MN and the myotube compartment for 24 h, at day 21 when the chemotactic and volumetric gradient was imposed (fig. 5f). Interestingly, TubA treatment had a striking effect on the neurite outgrowth in the P525L mutant FUS co-cultures, improving the outgrowth of neurites to a similar level as the isogenic control system (P525P, fig. 5g-i). TubA treatment also increased the outgrowth of neurites for the other *FUS* mutation (R521H) to a level above the control (supplementary fig. 9c-e), although it had no effect on the initial outgrowth of neurites during the first 24 h of culture (supplementary fig. 10).

Since TubA proved to have a beneficial effect on neurite outgrowth and HDAC6 recently has been demonstrated to regulate AChR receptor clustering (47), we next investigated whether this would have an effect on the formation of NMJs. Indeed, compared to our control NMJ data we observed that this short treatment with TubA already resulted in a tendency towards an increase in total and multiple contact point NMJ numbers in *FUS*-ALS P525L co-culture systems, although this did not reach statistical significance (fig. 5k, l). However, we could demonstrate a significant increase in multiple contact point NMJs in the *FUS*-ALS R521H co-cultures reaching similar levels as its corresponding control (R521R, fig. 5l).

In conclusion, we observe a beneficial effect of HDAC6 inhibition on neurite outgrowth and NMJ formation in mutant *FUS* co-cultures illustrating that our *in vitro* system can be used to test new therapeutic strategies increasing NMJ formation.

## Discussion

In this study, we developed a standardized co-culture model between hiPSC-derived MN and human primary MAB-derived myotubes in commercially available microfluidic devices resulting in the formation of functional human NMJs. We observed that supplementation of agrin and laminin promoted polarized motor neuron outgrowth as well as NMJ formation, further enhancing the output of this *in vitro* model. In contrast, mutations in *FUS* had a striking negative effect on both neurite outgrowth and NMJ numbers. We could successfully normalize these phenotypes by correcting the point mutations, emphasising that the impairments are a direct cause of the mutations in *FUS*. Last but not least, we discovered that HDAC6 inhibition can improve these neurite outgrowth impairments and can restore to a certain extent the formation of NMJs, further prompting HDAC6 inhibition as a potential therapeutic strategy in ALS.

A human NMJ model eliminates the challenge of interspecies variability and reduces the need for overexpression of mutated genes in animal models, which might not fully recapitulate human disease aetiology or pathology (58–60). In terms of NMJ morphology, it is evident that rodent NMJs are larger, more complex and less fragmented than human NMJs, and that human NMJs are more stable across human adult lifespan (61). These differences highlight the importance of a human model of NMJs for mechanistic insights in human NMJs physiology and disease. The model that we developed generates NMJ-like structures which resemble human NMJs from amputates (61). These similarities are seen in the morphological analysis of single contact point NMJs, which resemble “nummular” morphology described by Jones and colleagues (61). In addition, large, irregular multiple contact point NMJs are generated which appear to be unique to our system. These NMJs present a more complex morphology compared to the “nummular” NMJs and might therefore be a sign of further maturation. In multiple cases, we observed a continuation of the MN axons post-embedment into myotubes, which somewhat deviates from the distinct axon termination in endplate formation as seen *in vivo*.

An advantage of our system is the use of standardized stem cell technology in generating hiPSC-derived MNs. This method allows for complete adaptation to various MN diseases, and can therefore be used in studies of a broad range of neurodegenerative disorders including sporadic forms of ALS, which is otherwise difficult to model. In addition, the use of MABs in a NMJ co-culture model has to our knowledge not been reported before. MABs present a valid alternative to human satellite cells or myoblasts, since they age less, can generate more passages and are generally easier to keep in culture (39). The MABs are successfully differentiated into myotubes, which can be cultured up to 10 days from initial differentiation, and effectively generate NMJs when co-cultured with the hiPSC-derived MNs. The limited time-window in which MAB-derived myotubes can be cultured is due to the detachment of the fibers. Prolonging myotube viability *in vitro* and hence increased NMJ maturity could likely be accomplished by the incorporation of a dynamic myotube adherence system allowing for myotube contractility (37). However, this requires a redesign of the commercial microfluidic device. An ideal model would contain both iPSC-derived MNs and myotubes from the same donor, but no high-yield, reproducible protocol to generate hiPSC-derived myotubes is available (62). Current protocols either rely on selective overexpression of myogenic genes or mimicking embryonic development with paracrine supplementation of small molecules (62). We tested some of these protocols but they displayed large variability and were therefore not yet compatible with our standardized system (results not shown).

Electrophysiology using patch clamp analysis is considered the golden standard for assessing neuronal activity, but due to the closed compartmentalized system of the microfluidic device, this is not possible. Ideally, the combination of multi-electrode arrays (MEAs) and microfluidic systems will most likely contribute to the electrophysiological assessment of motor neuron and NMJ activity.

As an alternative, chemical depolarization successfully generated an intracellular response, which activated the NMJs and induced of Ca^2+^ waves in the myotubes. We observed a successful innervation and activation of approximately 68% of the active myotubes. As the field of vision is determined *at random*, it also considers myotubes, which are not innervated by MN axons. This means that the amount of active myotubes might vary depending on the area of analysis.

The fluidic isolation in the microfluidic device is an important advantage, as it can ensure compartmentalized treatment of either MNs or myotubes, which is favourable in drug target evaluations. Implementations of chemotactic and hydrostatic gradients likewise create physical stressors, which upon manipulation can modulate the co-culture in a desired manner allowing for large adaptability. By using devices with different microgroove lengths, it is likewise possible to assess the polarized growth of MN axons, and to study pathophysiological changes in proximal or distal regions of the axon (29,33). Furthermore, our model facilitates the investigation of synergies between muscle and MNs in an unbiased way.

The pre-patterning of AChR on the surface of muscles is recognized by branching neurites and facilitates guidance of the MN axons towards the muscle structure resulting in the formation of NMJs (43). The combined treatment with agrin and laminin showed increased AChR clustering *in vitro* (43–45,63,64), and agrin supplementation was successful in enhancing NMJ formation in a 3D one-compartment model using human embryonic stem cell-derived MNs and human primary myoblasts (49). In addition, laminin reduced the threshold concentration of agrin required to induce AChR clustering (44,45). We observed an increase in the clustering of AChRs as well as the total number of NMJs due to the treatment with agrin and laminin in comparison to untreated controls. The amount of multiple contact point NMJs was also increased, while SEM analysis demonstrated an enhanced neurite embedding and axonal branch fanning. Interestingly, agrin and laminin supplementation also improved the neurite outgrowth of MNs.

Using this co-culture model, we already investigated the effects of mutant *FUS* on the formation of human NMJs. Interestingly, we saw a dramatic reduction in total NMJs with an almost 50% decrease in the number of NMJs per myotube in the P525L mutant *FUS* system in comparison to its isogenic control (P525P). The FUS mutations did not affect the amount of single contact point NMJs, but had a large impact on the number of multiple contact point NMJs. These findings are in line with the ‘dying back’ mechanism where mature NMJs are lost in disease, while newly formed immature NMJs are made simultaneously to compensate for the lack of muscle innervation. However, we did not observe a significant increase in single contact point NMJs in the mutated system, which could explain such a compensatory mechanism although a tendency could be observed. As a consequence, we cannot exclude that the P525L findings are a result of a delayed maturation of the system influenced by the impairment in neurite outgrowth caused by the ALS mutation. Importantly, as we do not report a clear neurite outgrowth deficiency in the R521H motor unit, this indicates that NMJ numbers are affected independently of maturation of the system but rather due to the ALS-causing mutation itself. The smaller difference in NMJ numbers in the other *FUS*-ALS patient pair (R521H and R521R) correlates with a general observation of mild phenotypes in this late-onset R521H patient in comparison to rather aggressive phenotypes found in the early-onset P525L patient in this system. More and more studies show impaired neurite outgrowth in ALS. These include differences in axonal branching (65), impairments in neurite elongation speed in a sporadic ALS motor unit (37) and in neurite length in iPSC-derived MNs from both familial and sporadic ALS patients (66). However, the underlying mechanism has yet to be established. We observed that the mutant *FUS*-dependent reduction in neurite outgrowth could be rescued by the selective inhibition HDAC6. As a consequence, there could be a link between neurite outgrowth impairment and the widespread defects of the motor neuron axonal transport machinery, which has been observed in multiple ALS cases, as well as in other neurodegenerative disorders (reviewed in 53). HDAC6 inhibition had also some beneficial effect on the NMJ morphology, which for the P525L motor unit system could be linked to the improvement in neurite outgrowth. When more neurites are able to interact with myotubes, it could allow for a further maturation of the NMJ structures, overall improving the morphology in the model. Our studies on the both genetically determined and chemotherapy-induced peripheral neuropathies already demonstrated a correlation between an improvement of axonal transport due to HDAC6 inhibition and the reinnervation of NMJs *in vivo* (51,53,54,68). Further studies are crucial to investigate the potential of HDAC6 inhibition in ALS, although our *in vivo* results in a FUS mouse model already indicate that also other HDACs might play an important role (69).

Overall, we established a versatile and relatively easy to use motor unit system to study the functionality of human NMJs in culture. This co-culture model could have broad applications and is perfectly suited to test specific therapeutic strategies focussing on reinnervation and/or the stabilisation of NMJs.

## Materials and methods

### Cell lines and reagents

Healthy human control iPSCs (SBAD2 line) generated from a 51-year-old Caucasian male donor were provided by the Stem Cell Institute Leuven (SCIL, Leuven, Belgium). This line is derived by StemBANCC and was previously provided to SCIL by Dr. P. Jennings (Innsbruck University, Austria) as part of the EU-ToxRisk consortium (70). In addition, two previously characterised *FUS*-mutant iPSC lines from a 17-year old male ALS patient carrying a *de novo* mutation (P525L) and a 71-year old female ALS patient (R521H) (41,42). The *FUS*-ALS lines were systematically compared to their corresponding CRIPSR/CAS9 gene-edited isogenic control (P525P and R521R) (41,42). Human control primary mesoangioblasts (MABs) were obtained from a healthy 58-year-old donor (SCIL, Leuven, Belgium). Written informed consent was obtained from the subjects who provided their samples for iPSC generation and MAB harvest. MAB isolation and characterisation was approved by the medical ethics committee of the University Hospital Leuven (n° S5732-ML11268). The generation of iPSCs from the healthy control was approved by the UK’s main research ethics committee as part of the StemBANCC project, while the use of iPSC in this study was approved by the ethics committee of KU Leuven. The use of *FUS* patient fibroblasts and generation of iPSCs was approved by the ethics committee of the University Hospital Leuven (n° S50354 and S63792). All cells were routinely tested for mycoplasma contamination with MycoAlert Mycoplasma Detection Kit (Lonza, Rockland, ME, USA, Cat N° LT07-318). Chemicals and cell culture reagents were purchased from Thermo Fisher Scientific (Waltham, MA, USA) unless specified otherwise.

### Preparation of microfluidic devices

Microfluidic devices (Xona™ Microfluidics, Temecula, CA, USA; Cat N° SND75 (microgroove length: 75 μm) and Cat N° XC150 (microgroove length: 150 μm)) and 7.8mil Aclar 33C sheets (Electron Microscopy Sciences, Hatfield, PA, USA, Cat N° 50425-25) were sterilized in 70% ethanol and left to air-dry in the laminar flow. Devices were placed individually in 10 cm petri dishes for easy handling. SND75 devices and Aclar sheets were coated separately using 100 μg/ml poly L-ornithine (PLO) (Sigma, St. Louis, MO USA, Cat N° P3655-100MG) in DPBS (Cat N° 14190250) prior to mounting of devices on the Aclar sheet. XC150 devices were coated directly using 100 μg/mL PLO in DPBS. Importantly, avoidance of air bubble formation was ensured during the coating of the channels by never removing liquid directly from channels.

All coated material was incubated at 37°C, 5% CO_2_ for 3 h and subsequently washed twice in DPBS and once in sterile water. SND75 devices and Aclar sheets were left in the laminar flow cabinet for complete drying (20-30 min) before mounting of devices onto Aclar sheets. Assembly was performed in the laminar flow under a microscope with the use of a forceps to ensure the complete alignment of channels, grooves and borders of the wells. Once assembled, devices were checked for leakage and coated with 20 μg/ml laminin (Sigma, Cat N° L2020-1MG) in Neurobasal medium (Cat N° 21103049). A volume difference was established between the two sides of the device to allow laminin coating to pass through the microgrooves and the devices were incubated overnight at 37°C in 5% CO_2_. Overnight incubation hardened the silicone SND75 devices and further sealed them onto the Aclar sheets. The following day, devices were carefully flushed once with DPBS before plating neural progenitor cells (NPCs).

### Differentiation of iPSCs into motor neuronal progenitor cells and seeding in microfluidic device

The differentiation protocol utilized is based on a modified version of the Maury et al. protocol (71) and has previously been described (41,42,72). In brief, iPSCs were harvested using collagenase type IV (Cat N° 10780004) and collected in Corning^®^ ultra-low attachment flasks (Sigma, Cat N° 734-4140) to facilitate embryoid body formation. Cells were maintained in neuronal medium (50% DMEM/F12 (Cat N° 11330032) and 50% Neurobasal medium (Cat N° 21103049) with 0.5% L-glutamine (Cat N° 25030-024), 1% penicillin/streptomycin (Cat N° 15070063), 1% N-2 supplement (Cat N° 17502-048), 2% B-27™ without vitamin A (Cat N° 12587-010), 0.5 μM ascorbic acid (Sigma, Cat N° A4403), and 0.1% β-mercaptoethanol (Cat N° 31350010)) supplemented with 5 μM Y-27632 (Merck Millipore, Burlington, MA, USA; Cat N° 688001), 0.2 μM LDN-193189 (Stemgent, Beltsville, MA, USA; Cat N° 04-0074-02), 40 μM SB431542 (Tocris Bioscience, Bristol, UK, Cat N° 1614) and 3 μM CHIR99021 (Tocris Bioscience, Cat N° 4423) for 2 days with medium changes every day (=day 0-1). The following day (=day 2), neuronal medium was supplemented with 0.1 μM retinoic acid (Sigma; Cat N° R2625) and 500 nM smoothened agonist (Merck Millipore; Cat N° 566660), which was refreshed at day four. On day 7, 10 ng/ml BDNF (Peprotech, Rocky Hill, NJ, USA, Cat N° 450-02B) and 10 ng/ml GDNF (Peprotech, Cat N° 450-10B) was additionally added to the day four medium. On day 9, 20 μM DAPT (Tocris Bioscience, Cat N° 2634) was additionally supplemented to day 7 medium. On day 10, embryoid bodies were dissociated into a single cell NPC suspension using 0.05% trypsin (Gibco, Gaithersburg, MA, USA, Cat N° 25300054) and cryopreserved. NPCs were seeded in the two wells and channel on one side of the microfluidic device at 125,000 cells in 30-50 μl day nine medium supplemented with 1% RevitaCell™ (Cat N° A2644501) per well (total of 250,000 NPCs/device). After seeding, devices were incubated for 5-10 min at 37°C and 5% CO_2_ to allow cell attachment to the surface before additional day nine medium was added to all 4 wells for a total of 200 μl/well. Subsequently, 5-6 ml sterile DPBS was added to the 10 cm petri dish around the devices to avoid medium evaporation. Seeding in both wells and channel facilitated a larger and more robust network of MNs, which was less likely to detach during medium changes.

### Derivation and maintenance of human mesoangioblasts

Human MABs were isolated and cultured as previously described (39,40). Briefly, upon receipt of skeletal muscle biopsy, the tissue was minced and incubated for 2 weeks on collagen from calf skin-coated (Cat N° 17104019) 6-cm dishes in growth medium: 15% FBS (Cat N° 10270106), 1% sodium pyruvate (Life Technologies, Carlsbad, CA, USA, Cat N° 11360-070), 1% non-essential amino acids (Cat N° 11140050), 1% L-glutamine (Cat N° 25030-024), and 0.5% penicillin-streptomycin (Cat N° 15070063), and 1% insulin transferrin selenium (Cat N° 41400045) in IMDM (Cat N° 12440053) supplemented with 5 ng/ml recombinant human basic fibroblast growth factor (bFGF) (Peprotech, Cat N° 100-18B). Medium was changed every 4 days. After 14 days, cells were fluorescent activated cell (FACS)-sorted for human alkaline phosphatase (R&D systems, Minneapolis, MN, USA, Cat N° MAB1448) and expanded further in T75 flasks (Sigma, Cat N° CLS3276) on collagen from calf skin (Cat N° 17104019) in growth medium. Cells were cryopreserved in knockout serum replacement (Cat N° 10828-028) with 10% DMSO (Sigma, Cat N° D2650-100ML), passaged, or seeded in devices when reaching 70% confluence. For passaging MABs were washed once with DPBS, followed by incubation with TrypLE express (Cat N° 12605010) for 3 min at 37°C in 5% CO_2_. After incubation, TrypLE was neutralized with growth medium, and cells were gently scraped, collected and centrifuged for 3 min at 0.3 RCF, before cells were counted and reseeded in a T75 or T175 flask or device. Passaging was performed 1-2 times a week for cell expansion until a maximum passage number of 13. Since physical contact between MABs initiates fusion and lowers the myogenic potential, a cell confluence of 70% was never exceeded.

### Co-culturing myotubes and motor neurons in microfluidic device

On day 11 of motor neuron differentiation, day 10 medium was refreshed on both sides of the device keeping an equal volume across microgrooves. On day 14, 200 μl neuronal medium supplemented with 10 ng/ml BDNF, 10 ng/ml GDNF and 20 μM DAPT was added to each well to initiate NPC differentiation into spinal MNs (sMNs). On day 16, 200 μl day 14 medium additionally supplemented with 10 ng/ml CNTF (Peprotech, Cat N° 450-13B) was added to each of the four wells. On day 17, MABs were dissociated using TrypLE and seeded in the two wells and channel opposite to the MNs in the microfluidic device at 100,000 cells in 30-50 μl growth medium per well (total of 200,000 MABs/device). After seeding, devices were incubated for 5-10 min at 37°C in 5% CO_2_ to allow cell attachment to the surface before additional growth medium was added to the two wells for a total of 200 μl/well. MABs were large cells with spherical morphology when dissociated in suspension. On day 18, MN compartments received neuronal medium supplemented with 10 ng/ml BDNF, GDNF and CNTF, while MABs differentiation into myotubes was initiated in the opposite compartments using MAB differentiation medium containing 2% horse serum (Cat N° 16050122) and 1% sodium pyruvate in DMEM/F12 supplemented with 0.01 μg/ml recombinant human agrin protein (R&D Systems, Cat N° 6624-AG-050). At day 21, a 300% chemotactic growth factor gradient and 200% volumetric gradient were established to facilitate polarized axonal growth through the microgrooves of the microfluidic device towards the myotube compartment. MN compartments received 100 μl/well neuronal medium without neurotrophic factor supplements, while myotube compartments received 200 μl/well neuronal medium supplemented with 30 ng/ml BDNF, GDNF and CNTF in addition to 20 μg/ml laminin and 0.01 μg/ml agrin. The growth factor and volume gradients including laminin and agrin supplements were kept at each medium change, which was performed every other day until day 28 of MN differentiation equivalent to 10 days of co-culturing MNs and myotubes.

### Tubastatin A treatment

*FUS*-ALS and isogenic control co-cultures in XC150 devices were treated for 24 h with 1 μM Tubastatin A (TubA) (Selleckchem, Houston, TX, USA, Cat N° S8049) in day 21 medium and combined with the start of the chemotactic and volumetric gradient. TubA was added to both the soma and the myotube compartment. Control co-culture devices were kept in parallel without TubA treatment. At day 28, devices were analysed for neurite outgrowth and NMJ were quantified.

### Immunocytochemistry

Xona™ XC150 devices were used for immunocytochemistry (ICC) analysis of NMJ formation and neurite outgrowth, while MN differentiation verification and myotube fusion were imaged on 13 mm #1.5 coverslips (VWR, Monroeville, PA, USA, Cat N° 631-0150P) and in 96-well black tissue culture plates (Perkin Elmer, Waltham, MA, USA, Cat N° 6005430). Cells were washed once with DPBS before fixation using 2-4% paraformaldehyde (Cat N° 28908) for 15-20 min at room temperature (RT). After fixation, cells were permeabilized for 20 min at RT with DPBS + 0.1% Triton X-100 (Sigma, Cat N° T8787-250ML) followed by blocking for 20 min at RT in 5% normal donkey serum (Sigma, Cat N° D9663-10ML) in 0.1% PBS-Triton X-100. Primary antibodies (supplementary table 1) were diluted in 0.1% PBS-Triton X-100 with 2% normal donkey serum and incubated with a volume gradient between MN/myotube compartments overnight at 4°C.

The following day, cells were carefully washed and incubated with secondary antibodies (supplementary table 2) diluted in 2% normal donkey serum in 0.1% PBS/Triton X-100 for 1h at RT in the dark with a volume gradient between compartments.

To label nuclei, cells were incubated with DAPI (NucBlue Live Cell Stain ReadyProbes reagent, Cat N° R37605) for 20 min and coverslips were mounted with Fluorescence Mounting Medium (Dako, Glostrup, Denmark, Cat N° S3023), while wells in the device were sealed with one drop of mounting medium per well before acquiring images. Coverslips, plates and devices were imaged using an inverted Leica SP8 DMI8 confocal microscope and quantifications were performed utilizing Image J 1.52b software.

For MN quantifications, a minimum of 100 cells were randomly selected based on positive DAPI staining. Five random images at 20x magnification were acquired from each biological replicate. For myotube fusion index, nuclei were counted using Image J particle analyser tool with a nucleus size between 50-700 μm^2^. Five random images at 10x magnification were acquired from each biological replicate from each condition. For AChR cluster quantifications, a minimum of 20 random field of visions (=581.8 μm^2^) per condition were selected and myotubes was determined based on positive myosin heavy chain (MyHC) marker. MyHC-positive cells containing multiple nuclei were selected as myotubes. For NMJ quantifications, each image field was selected based on α-bungarotoxin (Btx)-positive clustering, recorded in z-stack, and the number of co-localizations between synaptophysin (SYP) and Btx was counted per myotube.

### Neurite outgrowth quantifications

The co-cultures used for NMJ quantification were likewise used for neurite outgrowth quantifications. In addition, MNs cultured in devices without myotubes were used to assess a potential myotube influence on neurite outgrowth. Tile scan images of NEFH fluorescence were taken at 10x magnification in a 1024×1024 format using an inverted Leica SP8 DMI8 confocal microscope. Tile scans were merged automatically using Leica software auto merge and auto stitching with a smooth overlap and linear blending. Afterwards, images were cropped at the microgroove edge displaying solely the myotube compartment with crossing neurites. Neurites were identified and isolated using ilastik 1.3.3post1 Pixel Classification software with a sigma of 0.3-1.6 for color/intensity, edge and texture. The outlined neurites were exported as probabilities predictions. Each image was adjusted for a threshold between 25-70 % (see supplementary table 3 for exact values) and converted to an 8-bit mask using Image J 1.52p software. Using Image J software’s particle remover plugin, pixels between 0-10 μm^2^ were removed. To assess neurite outgrowth, total number of pixel intersections were quantified utilizing a custom-made Image J 1.52p software script. The script performs a linear Scholl analysis and automatically quantifies the amount of pixel crossings per line intersection similar to a previously published method (48). The distance between each line intersection is 50μm. Due to high neurite density and bundle formation at the exit of the microgrooves, we omitted the measurements at the first line at 50 μm and started measuring at 100μm distance from the microgrooves.

For MN-NPC neurite outgrowth quantifications in mono-cultures, day 10 NPCs were plated in 24-well plates (Greiner bio-one cellstar, Vilvoorde, Belgium, Cat N° 662160) and imaged for 24 h using an IncuCyte ZOOM device with the IncuCyte ZOOM 2016A software and a Nikon S Plan Fluor ELWD 20X/0.45 OFN22 DIC N1 objective (Essen BioScience). Nine images per condition were taken every two hours. Cells were treated with 1-3 μM TubA for 24 h with control conditions receiving no treatment. Neurite outgrowth was analysed using the IncuCyte NeuroTrack Phase Neurites software with a brightness segmentation mode, a segmentation adjustment of 1, a cleanup min cell width of 10 μm, a neurite sensitivity of 0.5 μm and neurite width of 1 μm. The experiment was performed in three independent replicates with two technical replicates in each.

### Scanning electron microscopy

For SEM, myotubes and motor neurons were cultured in Xona SND75 devices on Aclar sheets and fixed with 2.5% glutaraldehyde in 0.1 M Na-cacodylate buffer pH 7.2 for 2 h at RT. After three washing steps in the same buffer, the microfluidic devices were carefully removed and Aclar sheets were clipped to round discs of 18 mm diameter. Subsequently, the discs were incubated in 1% osmium tetroxide for 1 h, washed in milli-Q water and dehydrated in a graded ethanol series to 100% ethanol. The discs were then inserted in a coverslip-holder for critical point drying in a Leica CPD300 apparatus for two hours, in which they were kept submerged in 100% ethanol at all times. Finally, the dried discs were mounted on SEM support stubs with carbon-stickers and coated with 4 nm Chromium in a Leica ACE600 coating machine. Cells and myotubes were studied and imaged in a Zeiss Sigma SEM at an accelerating voltage of 5 kV and at a working distance of 7 mm.

### Calcium fluorescent imaging

Xona™ XC150 devices were used for live-cell functionality assessment of NMJs. On day 28 of MN differentiation, myotube compartments were incubated for 25 min at 37°C, 5% CO_2_ with 200 μl/well neuronal medium supplemented with 30 ng/ml BDNF, GDNF and CNTF and 5 μM Fluo-4 AM (Cat N° F14201), while MN compartments received a medium change with 200 μl/well neuronal medium without neurotrophic factors. Fluo-4 AM is a Ca^2+^ indicator, which exhibits an increase in fluorescence upon Ca^2+^ binding. After incubation, the medium was refreshed on both sides in order to re-establish the volume and growth factor gradient. MNs were stimulated with 50 mM potassium chloride (KCl) in neuronal medium and the consequent Fluo-4 fluorescence was recorded in the myotube compartment (10x magnification with 1 s intervals for a duration of 1 min per set). Each field of vision was selected with bright field based on the presence of myotubes in the myotube compartment. For each device, the motor neuron stimulation was performed twice with approximately 2 min pause in between, followed by a positive test of myotube activity with direct stimulation of myotubes with 50 mM KCl in myotube compartment. MN compartment was stimulated once to confirm fluidic isolation and absence of Fluo-4 AM flow back through microgrooves during incubation. In addition, myotubes were cultured without MNs in 24-well black ibitreat μ-plates (Ibidi, Planegg, Germany, Cat N° 82406) and tested for Ca^2+^ functionality. To ensure the specificity of our experiment, myotube compartments were treated with 19 μM of the AChR competitive antagonist tubocurarine hydrochloride pentahydrate (DTC) (Sigma, Cat N° T2379-100G) 10 min before analysis. All recordings were acquired and analysed with a Nikon A1R confocal microscope and NIS-Elements AR 4.30.02 software. For functionality quantifications, each myotube was manually circled utilising NIS-elements Time Measurement tool and individually analysed for increase in Fluo-4 fluorescent signal over a 1-min time period. Increase in Ca^2+^ intensity was calculated as difference between peak value within first 30 recorded seconds and baseline. Baseline was calculated as average value of the first 5 seconds of recordings before KCl stimulation.

### Statistics

A biological replicate represents an independent MN differentiation from day 0 to day 28, and independent MAB differentiation into myotubes or an independent co-culture in a device. MN, myotube fusion index and AChR cluster quantifications were performed in three biological replicates, and NMJ quantifications in a total of four replicates with two technical replicates in each. NMJ functionality experiments were performed in four independent co-culture replicates and each experiment was performed in three technical replicates. NMJ blocking experiments with DTC were performed in three replicates. Neurite outgrowth quantifications were performed in three-four replicates using the healthy control iPSC line. For the *FUS*-iPSC lines, NMJ and neurite outgrowth quantifications were performed in three-eight replicates, and for TubA treatment in three-five replicates. Data are presented as mean ± s.e.m., unless indicated otherwise. Statistical analyses were made in GraphPad Prism 7.04. Data were tested for normal Gaussian distribution using Anderson-Darling test, D’Agostino-Pearson omnibus normality test and Shapiro-Wilk normality test. Statistical analyses were determined using unpaired t-test or Mann-Whitney test for differences of mean between two groups and one-way Anova with Tukey’s multiple comparisons test and Kruskal-Wallis test with Dunn’s multiple comparisons test for difference of mean between multiple groups. *p< 0.05, **p <0.01, ***p<0.001 and ****p <0.0001.

### Availability of data and materials

The data sets used and analysed during the current study are available from the corresponding author on reasonable request. The script used for neurite outgrowth quantifications is available from the corresponding author on request.

## Acknowledgements

The authors thank Sebastian Munck and Nikky Corthout from LiMoNe, Research Group Molecular Neurobiology (VIB-KU Leuven) for discussions concerning Ca^2+^ live-cell imaging experiments. This research was supported by the Fulbright Commission to Belgium and Luxembourg, the VIB, the KU Leuven (C1 and “Opening the Future” Fund), the “Fund for Scientific Research Flanders” (FWO-Vlaanderen), the Agency for Innovation by Science and Technology (IWT; SBO-iPSCAF), the Belgian Government (Interuniversity Attraction Poles Programme P7/16 initiated by the Belgian Federal Science Policy Office), the Thierry Latran Foundation, the “Association Belge contre les Maladies neuro-Musculaires” (ABMM) and the ALS Liga België (A Cure for ALS). TV is supported by a strategic basic research Ph.D. Grant awarded by the FWO (1S60116N). PVD holds a senior clinical investigatorship of FWO-Vlaanderen and is supported through the E. von Behring Chair for Neuromuscular and Neurodegenerative Disorders and the KU Leuven funds “Een Hart voor ALS”, “Laeversfonds voor ALS Onderzoek” and the “Valéry Perrier Race against ALS Fund”.

## Authors’ contributions

KSD, LF and TV planned and designed the experiments. ENK and KSD optimised the system. KSD performed most of the experiments and data analysis. LF and KSD performed confocal microscopy. KSD and PB performed scanning electron microscopy experiments. AK assisted in neurite outgrowth quantifications. BP wrote the Image J neurite outgrowth script. MS provided mesoangioblasts and valuable ideas for the project. GG provided advice for mesoangioblast culturing and differentiation. PVD provided ideas for the project. LVDB planned and supervised the project. KSD wrote the paper. LF, TV and LVDB edited the manuscript. All authors read and approved the final version of the paper.

