## Supplementary Material for "Human motor units in microfluidic devices are impaired by FUS mutations and improved by HDAC6 inhibition"

### Supplementary information

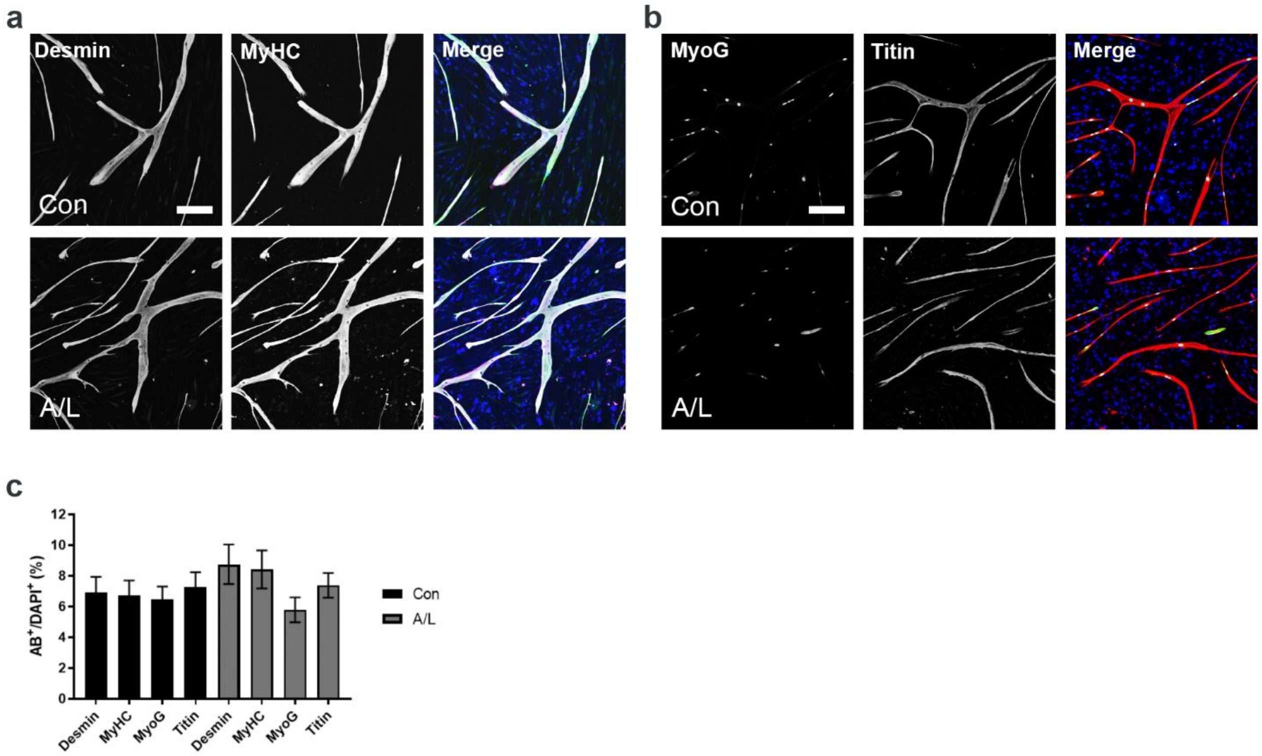


**Supplementary fig. 1: Myotube fusion index**

**a-b.** Representative confocal images of myotube markers: desmin, myosin heavy chain (MyHC), myogenin (MyoG) and titin in agrin (0.01 µg/ml) and laminin (20 µg/ml) supplemented conditions (A/L) and untreated controls (Con) after 10 days of differentiation. **c**. Quantification of myotube fusion index (n=15). Control data are identical to fig. 1 panel d. All data represent mean ± s.e.m. from three independent experiments and statistical analysis in panel **c** was performed using one-way Anova and Tukey's multiple comparisons test.

a b

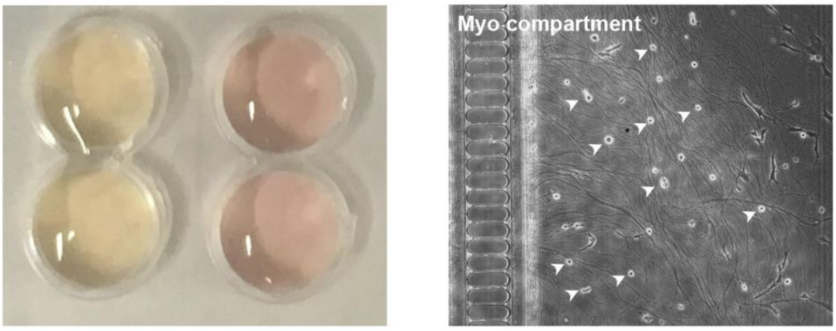


**Supplementary fig. 2: Fluidic isolation in microfluidic device and optimization of co-culture protocol**

**a.** Confirmation of fluidic isolation between left and right compartments in an XC150 microfluidic device. By establishing a volumetric gradient, the pH-sensitive medium on the motor neurons changes colour to yellow (left side), whereas the medium on the non-seeded compartment remains pink (right side) after 24 h. **b.** Maturation of MN in a device for 2 weeks (=day 24) before seeding of MABs (arrowheads). In order to prolong MN maturation and the sustainability of the co-culture system, we attempted to plate MABs at day 24 of MN differentiation. However, two weeks of MN maturation in the device resulted in large amounts of spontaneous neurite crossing, which inhibited the attachment of MABs in the channel (arrowheads). Due to the lack of myotube formation, we performed the seeding of MABs at day 17 (Fig 1.e).

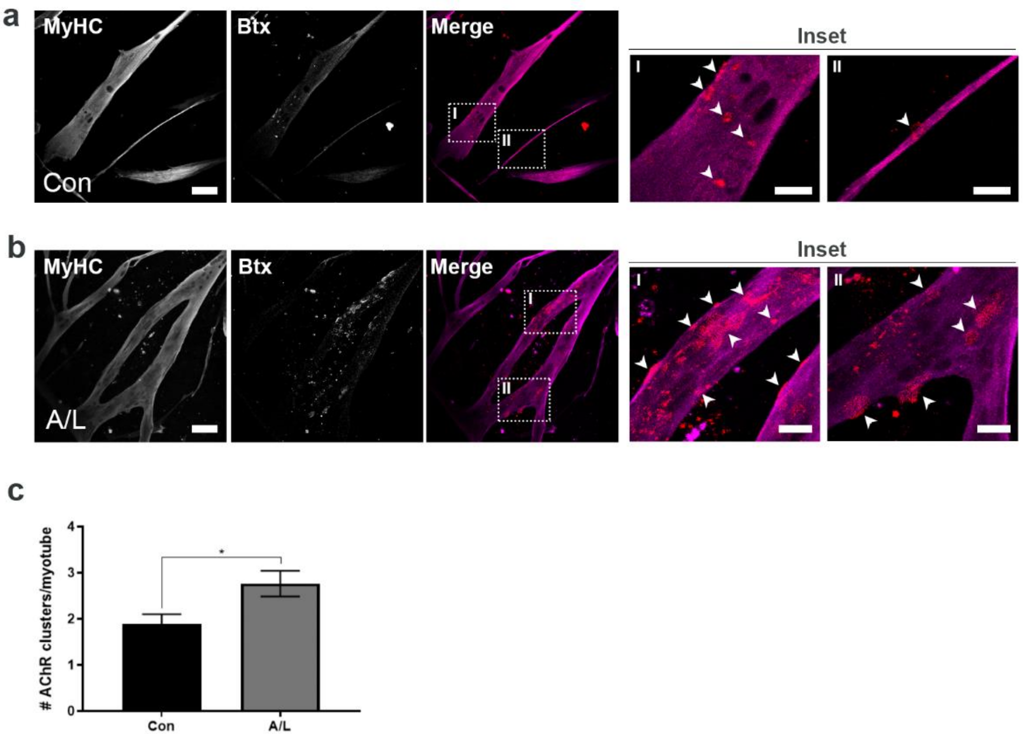


**Supplementary fig. 3: Agrin and laminin treatment enhance acetylcholine receptor clustering**

**a-b.** Representative ICC images of MyHC-positive myotubes with Btx-positive AChR clusters after 10 days differentiation in control conditions (Con) and with supplements of agrin and laminin (A/L). Scale bar: 75 µm. Insets represent a magnification of AChR clusters (arrowheads). Inset scale bar: 25 µm. **c.** Quantifications of AChR cluster number per myotube (n=20-29). The data represent mean ± s.e.m. and statistical analysis was performed using unpaired t-test with *$p<0.05$.

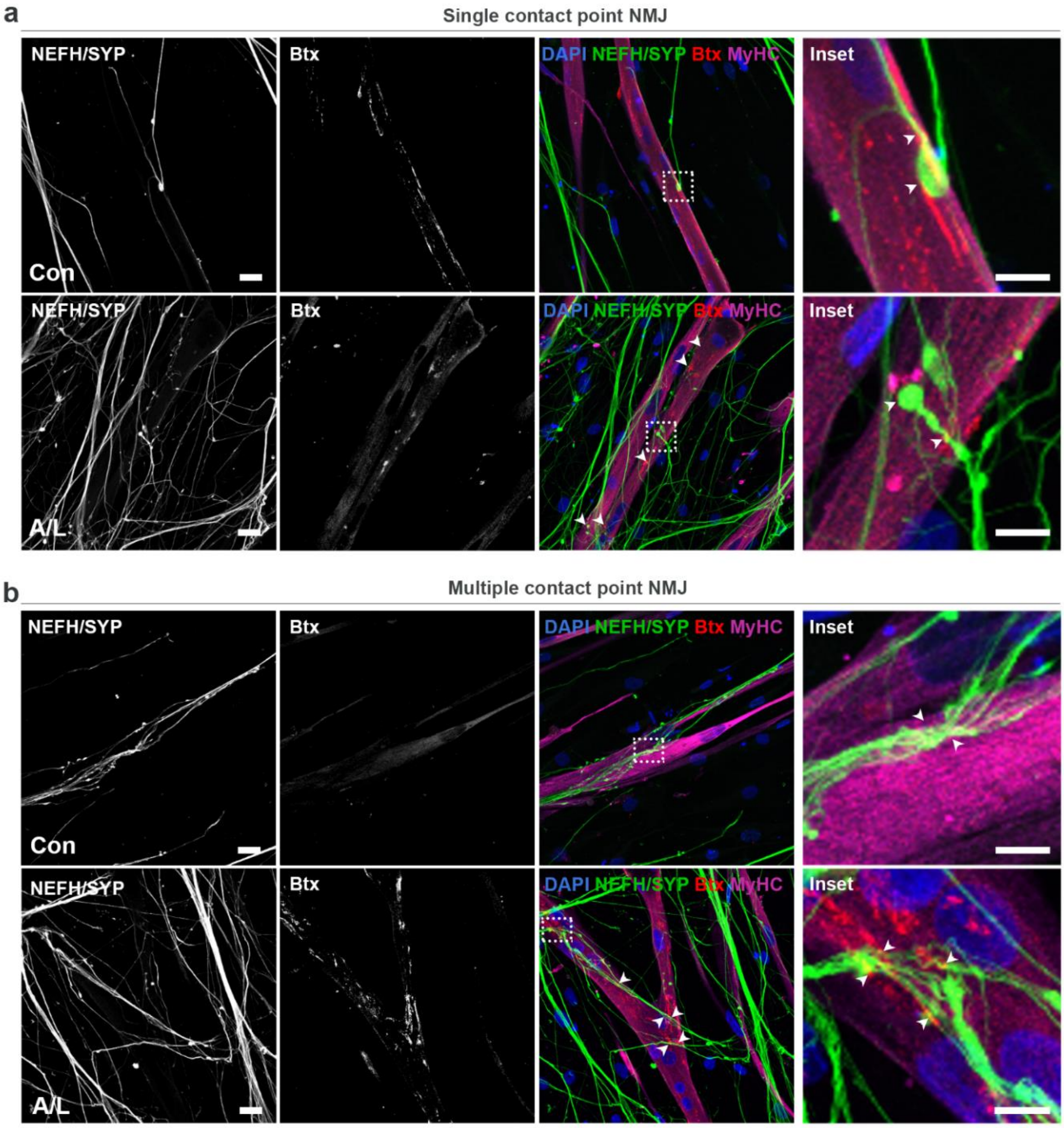


**Supplementary fig. 4: NMJ images related to fig. 3**

**a-b.** Confocal micrographs of NMJs in agrin and laminin supplemented conditions (A/L) and untreated controls (Con) at 28 days of motor neuron differentiation in XC150 microfluidic devices. NEFH- and SYP-positive motor neuronal axons either form a single contact point connection with myotubes forming rudimental NMJs (**a**) or they fan out upon interaction with myotubes and create multiple contact point NMJs (**b**). Scale bar: 25 µm. Arrowheads mark co-localizations between SYP and Btx. Inset scale bar: 10 µm.

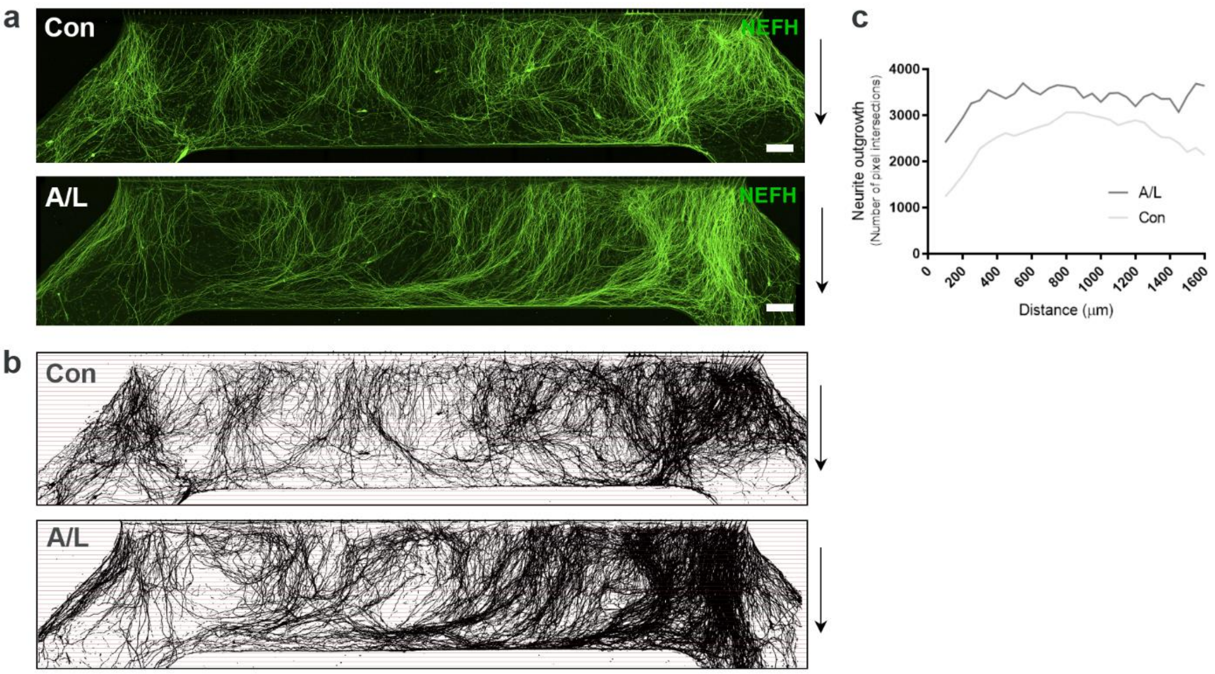


**Supplementary fig. 5: Agrin and laminin improve neurite outgrowth**

**a.** Tile scan confocal overviews of neurite outgrowth in myotube compartment. Arrows (right) depict growth direction from exit of microgrooves. Scale bar: 300 µm. **b.** Masks of tile scans with intersection lines at every 50 µm from microgroove exit. **c.** Neurite outgrowth quantifications of the number of pixel intersections in each condition (A/L and Con). Panel **c** was performed in three-four independent experiments with mean graphs shown.

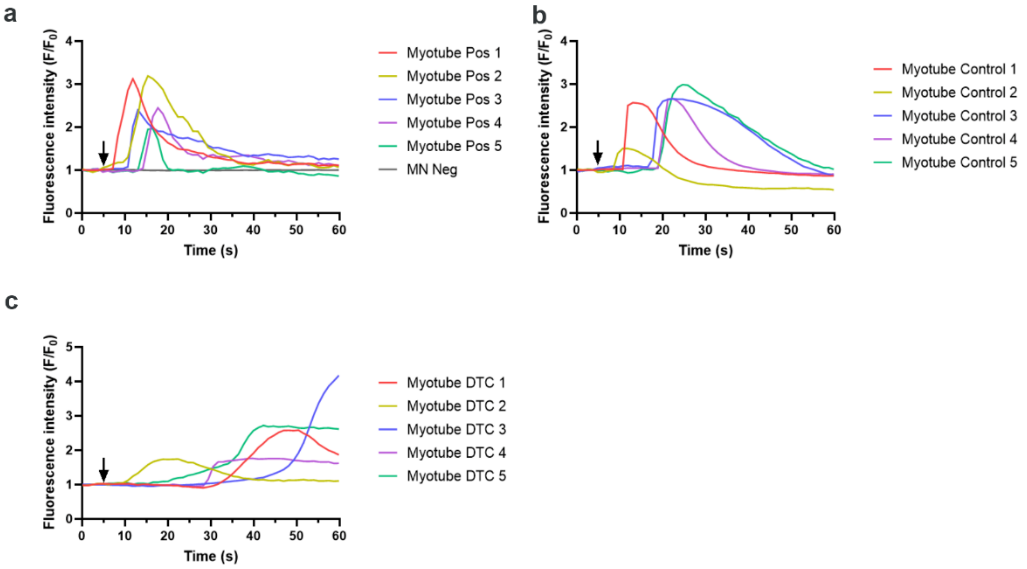


**Supplementary fig. 6: Myotube functionality controls related to fig. 4**

**a**. Representative $Ca^{2+}$ influx curves in myotubes in co-culture with MNs after direct KCl stimulation (arrow) of myotubes in myotube compartment (Myotube Pos 1-5). MN Neg curve represent a recording of the MN compartment after MN stimulation with KCl confirming a fluidic isolation between compartments during imaging. **b**. Representative $Ca^{2+}$ influx curves in myotubes cultured without MNs. Arrow mark KCl stimulation. **c**. Representative $Ca^{2+}$ influx curves in myotubes cultured without MNs after 10 min treatment with 19 µM nicotinic AChR competitive antagonist tubocurarine. Arrow mark KCl stimulation. Data represents three-four independent experiments.

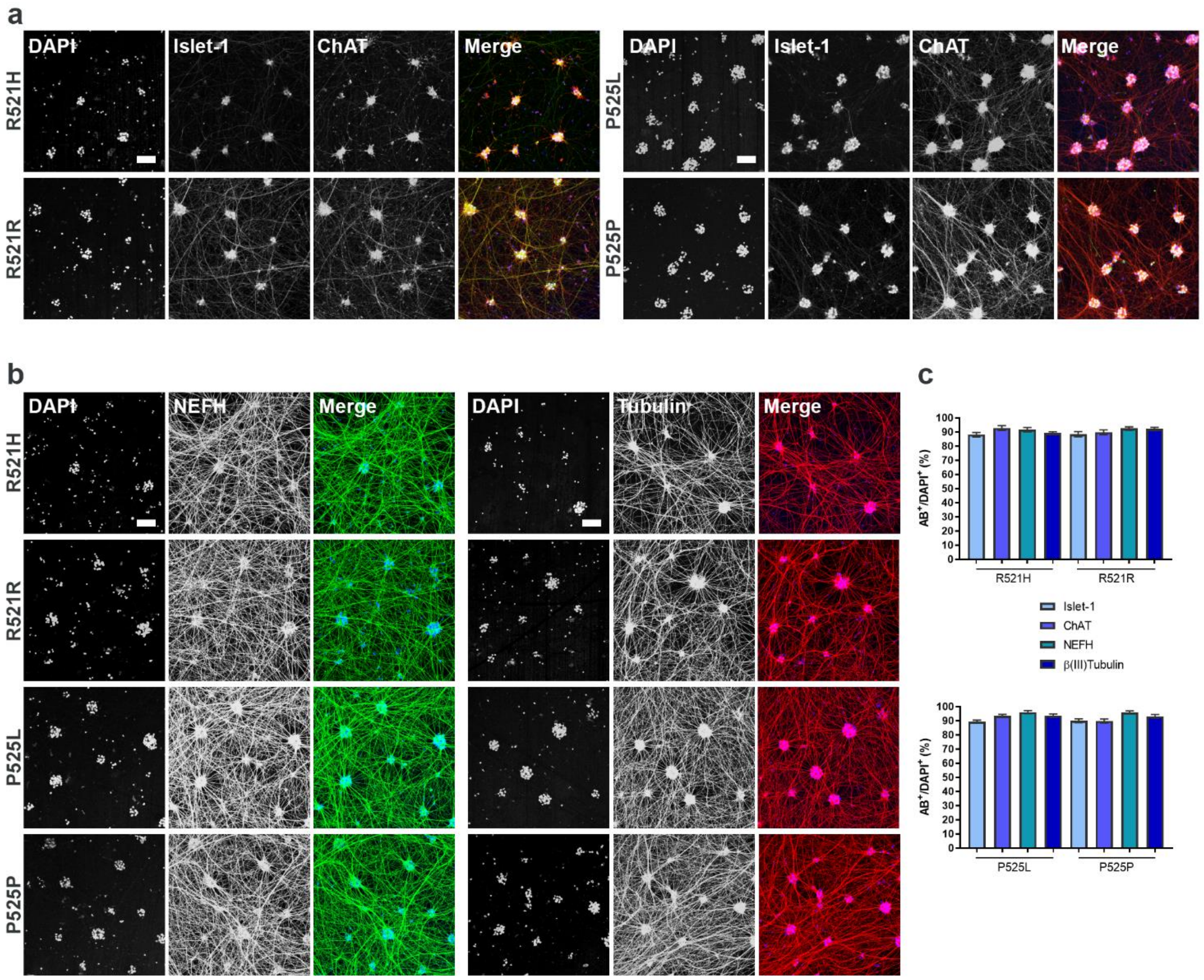


**Supplementary fig. 7: *FUS*-ALS MN differentiation verification**

**a-b.** Confocal images of mutant *FUS* MNs (P525L, R521H) and isogenic control MNs (P525P, R521R). MNs are stained with MN markers Islet-1 and choline acetyltransferase (ChAT) (**a**) and neurofilament heavy chain (NEFH) as well as the pan-neuronal marker βIII-tubulin (**b**) at day 28 of MN differentiation. Scale bar: 75 µm. **c.** Relative number of cells positive for MN and pan-neuronal markers (n=15). The data in panel **c** represent mean ± s.e.m and statistical analysis was performed using Kruskal-Wallis test with Dunn's multiple comparisons test from three independent experiments.

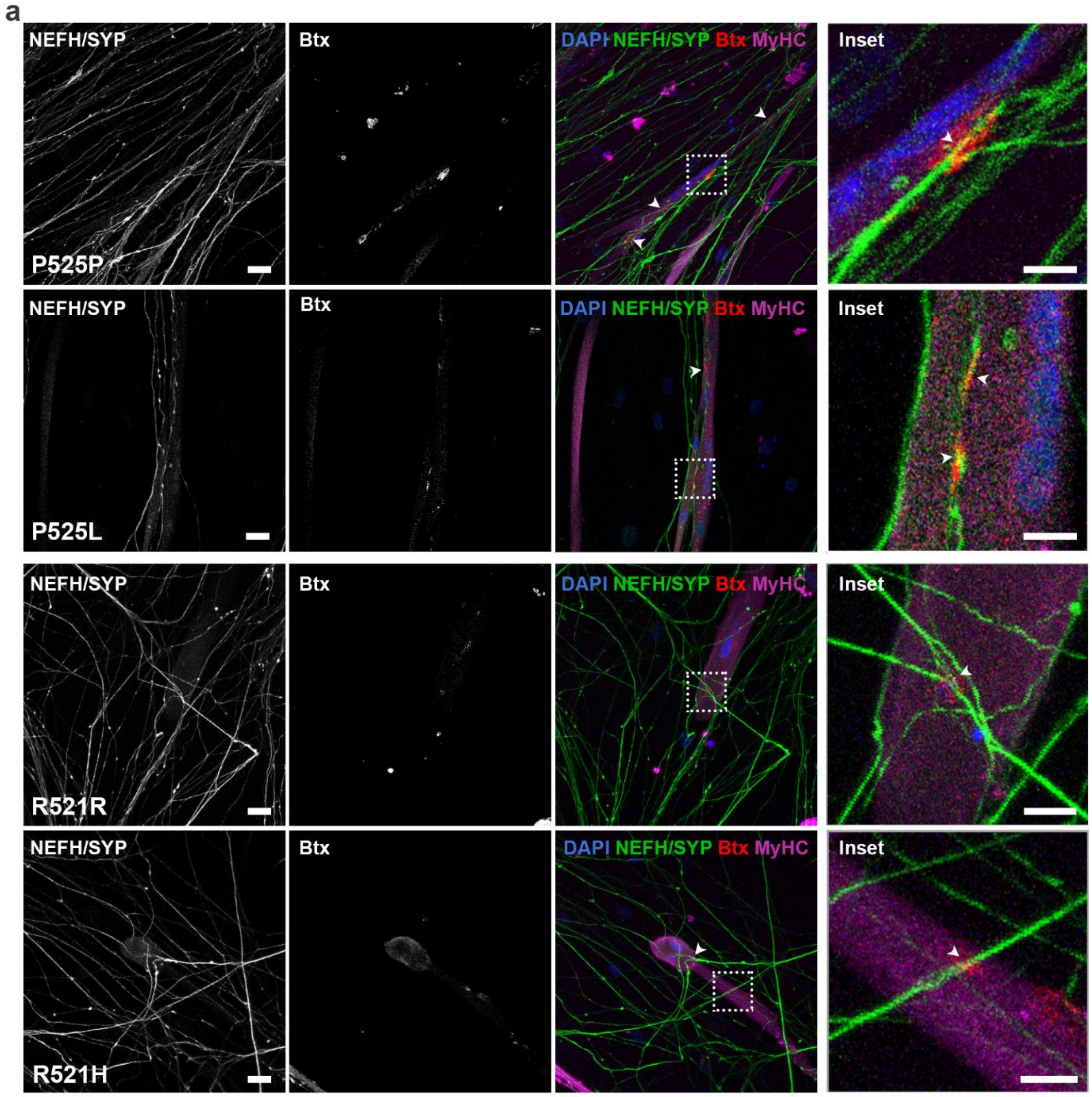


**Supplementary fig. 8: NMJ images related to fig. 5**

**a**. Confocal micrographs of NMJs in agrin and laminin supplemented conditions from *FUS*-mutant MN/myotube co-cultures (P525L, R521H) and isogenic control MN/myotube co-cultures (P525P, R521R) at 28 days of motor neuron differentiation in XC150 microfluidic devices. Scale bar: 25 µm. Arrowheads mark co-localizations between SYP and Btx. Inset scale bar: 10 µm.

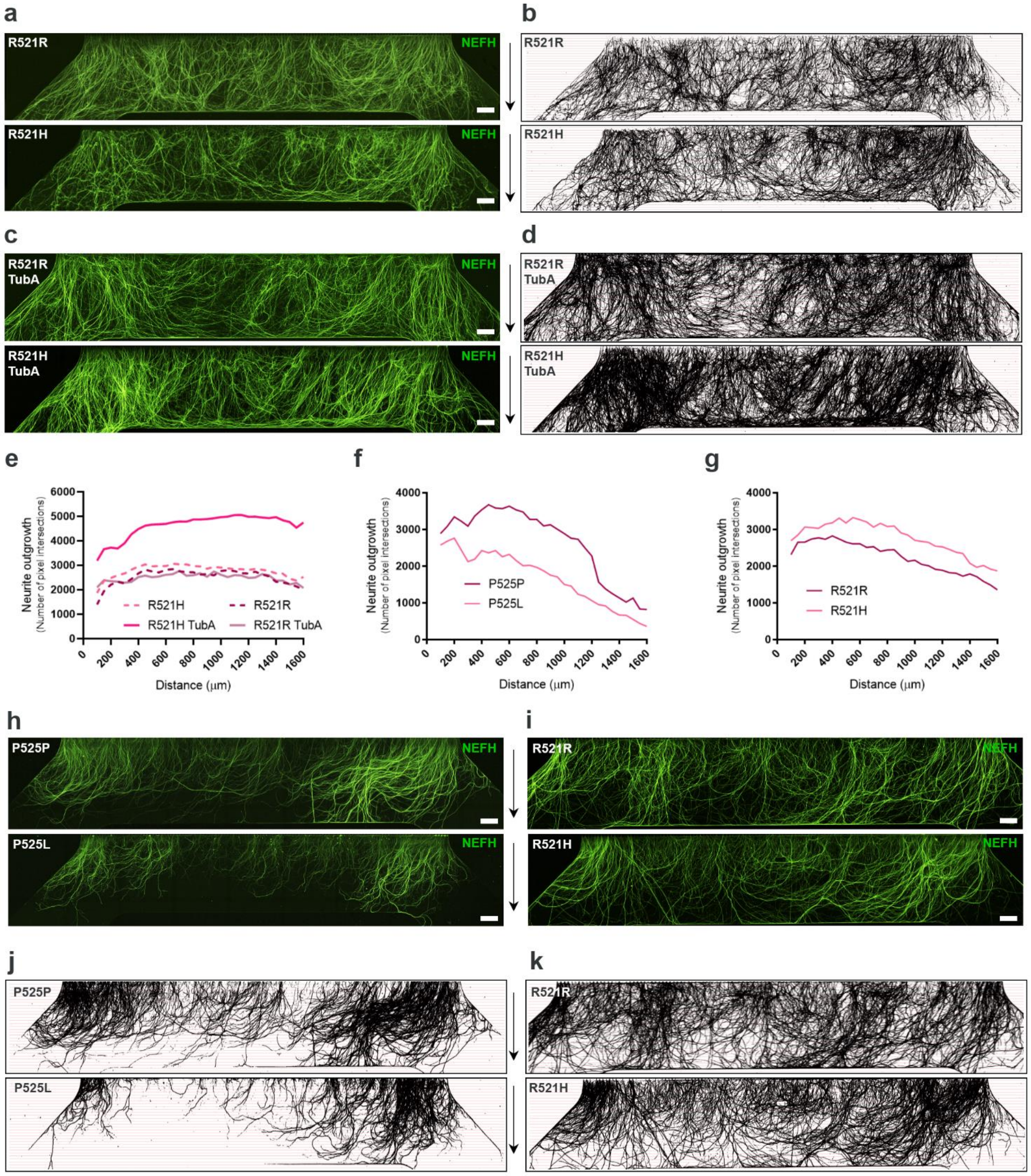


**Supplementary fig. 9: Neurite outgrowth related to fig. 5**

a. Tile scan confocal overviews of neurite outgrowth (NEFH) in myotube compartment from mutant *FUS* (R521H) and its isogenic control (R521R) MN/myotubes co-cultures. Arrows (right) depict growth direction from exit of microgrooves. Scale bar: 300 µm. b. Masks of tile scans with intersection lines at every 50 µm from microgroove exit. c. Tile scan images of neurite outgrowth in R521R and R521H MN/myotube co-cultures after 24 h of Tubastatin A (TubA) treatment. Scale bar: 300 µm. d. Masks of tile scans after TubA treatment. e. Neurite outgrowth quantifications of pixel intersections in R521R and R521H MN/myotube co-cultures before and after treatment with TubA. f-g. Neurite outgrowth quantifications in myotube compartment from mutant *FUS* (P525L, R521H) and corresponding isogenic controls (P525P, R521R) of the number of pixel intersections in MN cultures without myotubes. h-i. Tile scan confocal overviews of neurite outgrowth in MN cultures without myotubes. j-k. Masks of tile scans cultured without myotubes with intersection lines at every 50 µm from microgroove exit. All data are from three to eight independent experiments.

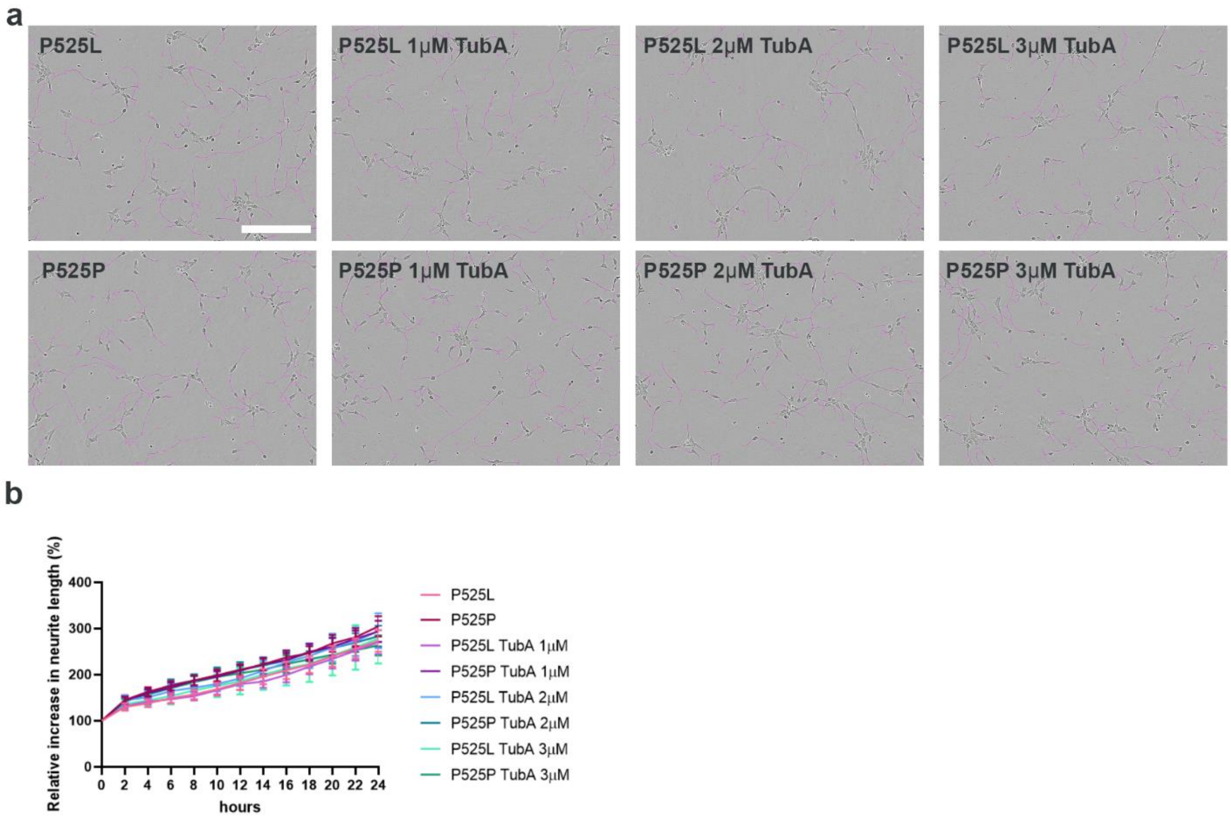


**Supplementary fig. 10: Neurite outgrowth in mono-culture is unaffected by *FUS*-ALS and HDAC6 inhibition**

**a**. Representative IncuCyte bright field images of *FUS*-ALS patient line P525L and isogenic control P525P with 1-3 µM Tubastatin A (TubA) treatment and controls receiving no treatment. Neurite are marked using IncuCyte Phase Neurite mask. Scale bar: 200 µm. **b**. Quantifications of relative increase in neurite length. Data represent mean ± s.e.m and statistical analysis was performed using one-way Anova with Tukey’s multiple comparisons test from three independent experiments.

**Supplementary table 1: Primary antibody overview**

| Antibody | Dilution | Company | Antibody Registry* Identifier |
|---|---|---|---|
| Rabbit anti-neurofilament heavy chain (NEFH) | 1:1000 | Abcam, Cat N° AB8135 | AB_306298 |
| Rabbit anti-synaptophysin (SYP) | 1:1000 | Cell Signaling, Cat N° 5461S | AB_10698743 |
| Mouse anti-myosin heavy chain (MyHC) | 1:20 | In-house, SCIL | |
| Goat anti-choline acetyltransferase (ChAT) | 1:500 | Millipore, Cat N° ab144P | AB_2079751 |
| Rabbit anti-islet 1 | 1:400 | Millipore, Cat N° ab4326 | AB_10563961 |
| Mouse anti-βIII-tubulin | 1:500 | Abcam, Cat N° ab7751 | AB_306045 |
| Mouse anti-titin | 1:300 | Developmental Studies Hybridoma Bank, Cat N° 9 D10 | AB_528491 |
| Rabbit anti-myogenin (MyoG) | 1:500 | Abcam, Cat N° Ab124800 | AB_10971849 |
| Rabbit anti-desmin | 1:200 | Abcam, Cat N° Ab15200 | AB_301744 |

* https://antibodyregistry.org/

**Supplementary table 2: Secondary antibody overview**

| Antibody | Dilution | Company | Antibody Registry* Identifier |
|---|---|---|---|
| Alexa Flour™ IgG (H+L) 488 donkey-anti-rabbit | 1:1000 | Thermo Fisher Scientific, Cat N° A21206 | AB_2535792 |
| Alexa Flour™ IgG (H+L) 555 donkey-anti-goat | 1:1000 | Thermo Fisher Scientific, Cat N° A21432 | AB_2535853 |
| Alexa Flour™ IgG (H+L) 555 donkey-anti-mouse | 1:1000 | Thermo Fisher Scientific, Cat N° A31570 | AB_2536180 |
| Alexa Flour™ IgG (H+L) 647 donkey-anti-mouse | 1:1000 | Thermo Fisher Scientific, Cat N° A31571 | AB_162542 |
| α-bungarotoxin (Btx) Alexa Flour™ 555 | 1:1000 | Thermo Fisher Scientific, Cat N° B35451 | AB_2617152 |

* https://antibodyregistry.org/

#### Supplementary table 3: Neurite outgrowth thresholds

| **Sample ID (MN/Myotube co-cultures)** | **Threshold (%)** | **Sample ID (MN/Myotube co-cultures)** | **Threshold (%)** | **Sample ID TubA treatment (MN/Myotube co-cultures)** | **Threshold (%)** |
|---|---|---|---|---|---|
| A/L biological replicate 1 | 60 | R521H biological replicate 1 | 35 | P525L biological replicate 1 | 50 |
| A/L biological replicate 2 | 60 | R521H biological replicate 2 | 70 | P525L biological replicate 2 | 60 |
| A/L biological replicate 3 | 70 | R521H biological replicate 3 | 70 | P525L biological replicate 3 | 55 |
| A/L biological replicate 4 | 70 | R521H biological replicate 4 | 30 | P525P biological replicate 1 | 55 |
| Control biological replicate 1 | 40 | R521H biological replicate 5 | 50 | P525P biological replicate 2 | 50 |
| Control biological replicate 2 | 45 | R521H biological replicate 6 | 50 | P525P biological replicate 3 | 55 |
| Control biological replicate 3 | 60 | R521H biological replicate 7 | 50 | R521H biological replicate 1 | 60 |
| P525L biological replicate 1 | 40 | R521H biological replicate 8 | 50 | R521H biological replicate 2 | 50 |
| P525L biological replicate 2 | 40 | R521R biological replicate 1 | 50 | R521H biological replicate 3 | 50 |
| P525L biological replicate 3 | 45 | R521R biological replicate 2 | 60 | R521H biological replicate 4 | 60 |
| P525L biological replicate 4 | 50 | R521R biological replicate 3 | 50 | R521H biological replicate 5 | 50 |
| P525L biological replicate 5 | 45 | R521R biological replicate 4 | 45 | R521R biological replicate 1 | 30 |
| P525L biological replicate 6 | 50 | R521R biological replicate 5 | 35 | R521R biological replicate 2 | 25 |
| P525P biological replicate 1 | 70 | R521R biological replicate 6 | 50 | R521R biological replicate 3 | 50 |
| P525P biological replicate 2 | 50 | **Sample ID (MN without myotubes)** | **Threshold (%)** | **Sample ID (MN without myotubes)** | **Threshold (%)** |
| P525P biological replicate 3 | 60 | P525L biological replicate 1 | 35 | R521H biological replicate 1 | 50 |
| P525P biological replicate 4 | 60 | P525L biological replicate 2 | 35 | R521H biological replicate 2 | 40 |
| P525P biological replicate 5 | 50 | P525L biological replicate 3 | 35 | R521H biological replicate 3 | 30 |
| P525P biological replicate 6 | 35 | P525P biological replicate 1 | 35 | R521R biological replicate 1 | 50 |
| P525P biological replicate 7 | 55 | P525P biological replicate 2 | 35 | R521R biological replicate 2 | 50 |
| | | P525P biological replicate 3 | 35 | R521R biological replicate 3 | 40 |
